# Incomplete recombination suppression fuels extensive haplotype diversity in a butterfly color pattern supergene

**DOI:** 10.1101/2024.07.26.605145

**Authors:** Rishi De-Kayne, Ian J. Gordon, Reinier F. Terblanche, Steve Collins, Kennedy Saitoti Omufwoko, Dino J. Martins, Simon H. Martin

## Abstract

Supergenes can evolve when recombination-suppressing mechanisms like inversions promote co-inheritance of alleles at two or more polymorphic loci that affect a complex trait. Theory shows that such genetic architectures can be favoured under balancing selection or local adaptation in the face of gene flow, but they can also bring costs associated with reduced opportunities for recombination. These costs may in turn be offset by rare ‘gene flux’ between inverted and ancestral haplotypes, with a range of possible outcomes. We aimed to shed light on these processes by investigating the BC supergene, a large genomic region comprising multiple rearrangements associated with three distinct wing color morphs in *Danaus chrysippus*, a butterfly known as the African monarch, African queen and plain tiger. Using whole-genome resequencing data from 174 individuals, we first confirm the effects of BC on wing color pattern: background melanism is associated with SNPs in the promoter region of *yellow*, within an inverted subregion of the supergene, while forewing tip pattern is most likely associated with copy number variation in a separate subregion of the supergene. We then show that haplotype diversity within the supergene is surprisingly extensive: there are at least six divergent haplotype groups that experience suppressed recombination with respect to each other. Despite high divergence between these haplotype groups, we identify an unexpectedly large number of natural recombinant haplotypes. Several of the inferred crossovers occurred between adjacent inversion ‘modules’, while others occurred within inversions. Furthermore, we show that new haplotype groups have arisen through recombination between two pre-existing ones. Specifically, an allele for dark coloration in the promoter of *yellow* has recombined into distinct haplotype backgrounds on at least two separate occasions. Overall, our findings paint a picture of dynamic evolution of supergene haplotypes, fuelled by incomplete recombination suppression.

## Introduction

Genomic architectures that promote co-inheritance of certain combinations of alleles at multiple loci can be beneficial if they maintain adaptive allele combinations in the face of genetic mixing [1]. Such genetic architectures are referred to as supergenes, because they facilitate the inheritance of complex phenotypes in a simple Mendelian fashion, (reviewed in [2]). Supergenes are often associated with chromosomal inversions, which have the unique feature of reducing recombination between haplotypes with distinct inversion orientations while allowing free recombination between haplotypes with the same orientation. This effectively divides the population into two subpopulations over a defined portion of the genome [3]. Supergenes have been found to underpin trait polymorphisms under balancing selection, such as alternative life-history and reproductive strategies [4–8] and mimetic coloration [9]. Inversions can also be favoured during local adaptation in the face of gene flow between two distinct environments if they maintain locally adapted combinations that differentiate ecotypes [10–12]. Here we include these cases of locally adapted inversions under the umbrella term ‘supergenes’, provided they still act to maintain allele combinations that would otherwise be broken down through gene flow and recombination. Genomic studies of species that exhibit local adaptation frequently uncover inversions underpinning differences between ecotypes [13–16], and ‘bottom-up’ analyses have identified signatures consistent with locally adapted inversions even when the specific traits they may be associated with are unknown [17–19]. This suggests that supergene architectures may be ubiquitous in nature, highlighting a need to better understand their evolutionary dynamics.

Recent theoretical and empirical work suggests that supergenes may evolve over time in their structure, composition, and effects on phenotype and/or fitness. First, supergene structure can evolve through recurrent chromosomal rearrangements. Studies across multiple systems show that supergenes often comprise several rearrangements, implying stepwise growth in the region of suppressed recombination (e.g. in *Heliconius* butterflies [20], *Danaus* butterflies [21], and *Solenopsis* fire ants [6]). In addition to allowing the incorporation of additional co-adapted alleles at other loci, subsequent rearrangements of the supergene region can cause recombination suppression between more than two distinct haplotypes [21]. Second, even without physical expansion, the repertoire of traits affected by an inversion supergene may expand over time as additional alleles become established at other loci within the region of recombination suppression [11,17]. The fitness consequences of a supergene could also change if selective pressures change over time or across space, potentially limiting the value of a supergene in a changing environment or during dispersal into a new environment [22]. Even in a stable environment, inversion supergenes may be subject to accumulation of increased mutational load compared to the rest of the genome due to their reduced opportunities for recombination and reduced effective population size (*N_e_*) (due to the effective subdivision of the population in that part of the genome) [23–25]. Some theoretical models suggest that this may lead to heterokaryotype advantage through sheltering of recessive deleterious mutations (associative overdominance) [23–25], or failure of locally adapted inversions to reach high frequency [26].

Another process that contributes to the evolution of inversion supergenes is rare recombination in heterokaryotypes, which results in some degree of gene flux between haplotypes. While single crossovers within inversions are usually strongly suppressed due to the production of unbalanced chromosomes [27], large inversions may allow for double crossovers [28], which result in the exchange of genetic material between arrangements, effectively creating a mosaic haplotype. In addition, gene flux of short fragments can occur through non-crossover gene conversion. This has been shown experimentally in *Drosophila* to occur at least as often within inversion heterokaryotypes as in syntenic regions [28–33]. Indirect inference in natural populations of a range of different species support the existence of considerable gene flux within inversions [32,34–40], and suggest that a drift-flux equilibrium can be reached [3]. Gene flux reduces linkage disequilibrium within inversions [41] and provides a means by which they may avoid some of the costs of recombination suppression described above, potentially facilitating their long-term persistence [6,23,37]. In some supergenes, sequence divergence suggests persistence of polymorphism for millions of years, even through multiple speciation events [21,37]. However, the role of gene flux in supergene temporal dynamics remains under-explored. A compelling example of recombination creating a novel haplotype associated with a distinct phenotypic class is seen in the supergene that controls reproductive morphology and behaviour in the ruff [5,37]. On the other hand, a mimicry supergene in *Papilio* butterflies shows phylogenetic relationships consistent with wholesale ‘allelic turnover’, in which new haplotypes arise (possibly via recombination) and replace ancestral ones [42]. Case studies such as these provide valuable insights into the range of processes contributing to supergene evolution.

*Danaus chrysippus*, a butterfly known as the African monarch, African queen, and plain tiger, presents an opportunity to investigate the contributions of the above processes to the evolution of a large, ancient supergene. Like other milkweed butterflies, *D. chrysippus* has bright warning patterns that advertise its toxicity. Across its range, it is divided into several parapatric morphs with distinct warning patterns that meet in a broad hybrid zone in eastern central Africa (Fig. 1A) [43,44]. Phenotypic differences in two forewing traits are controlled by the ‘BC supergene’ on chromosome 15, which links at least two color patterning loci, that were originally described in crossing experiments: The ‘B’ locus affects whether background color is dark/brown (dominant *B* allele) or pale/orange (recessive *b* allele) and the ‘C’ locus determines whether the apical black tip and white band is present (recessive *c* allele), or absent (dominant *C* allele) [21,45–48]. Previous genomic comparisons of a limited number of populations revealed the existence of three divergent haplotype groups (also called ‘alleles’) at the BC supergene, which were named according to the morphs in which they are found: ‘orientis’ (*Bc*, Southern Africa), ‘klugii’ (*bC*, East Africa) and ‘chrysippus’ (*bc*, West Africa, North Africa, Mediterranean, Asia) [45]. Note that while a distinct morph ‘alcippus’ is found in West Africa, it only differs in its hindwing phenotype, controlled by a locus on a different chromosome, but shares its forewing phenotype and corresponding *bc* haplotype with the chrysippus morph. Chromosome-scale assemblies of each of the three haplotypes revealed that, rather than comprising a single large inversion, the BC supergene is underpinned by a complex, modular rearrangement including several inversions and a large copy-number variable (CNV) region [21]. This complex architecture helps to explain how recombination is suppressed between more than two divergent haplotype groups. Phylogenetic analysis showed that the rearrangements probably occurred in a stepwise manner, beginning several million years ago, and that the supergene has persisted through multiple speciation events [21]. Intriguingly, an additional rearrangement has recently occurred and risen to high frequency in a small region of East Africa: a fusion of chr15 to the female-specific W chromosome [45,49]. Because crossing-over is limited to males in Lepidoptera, this fusion represents an additional mechanism of recombination suppression, effectively creating a new female-specific sub-group of the chrysippus haplotype [45]. There is also evidence for rare recombination within the supergene based on phenotypes in crossing experiments, and a putative recombinant haplotype assembly from a hybrid-zone individual [21,50]. Taken together, the above findings suggest that, despite its age, the BC supergene continues to evolve, raising questions about the full extent of diversity at this locus and the role of recombination in shaping this diversity.

**Figure 1.**
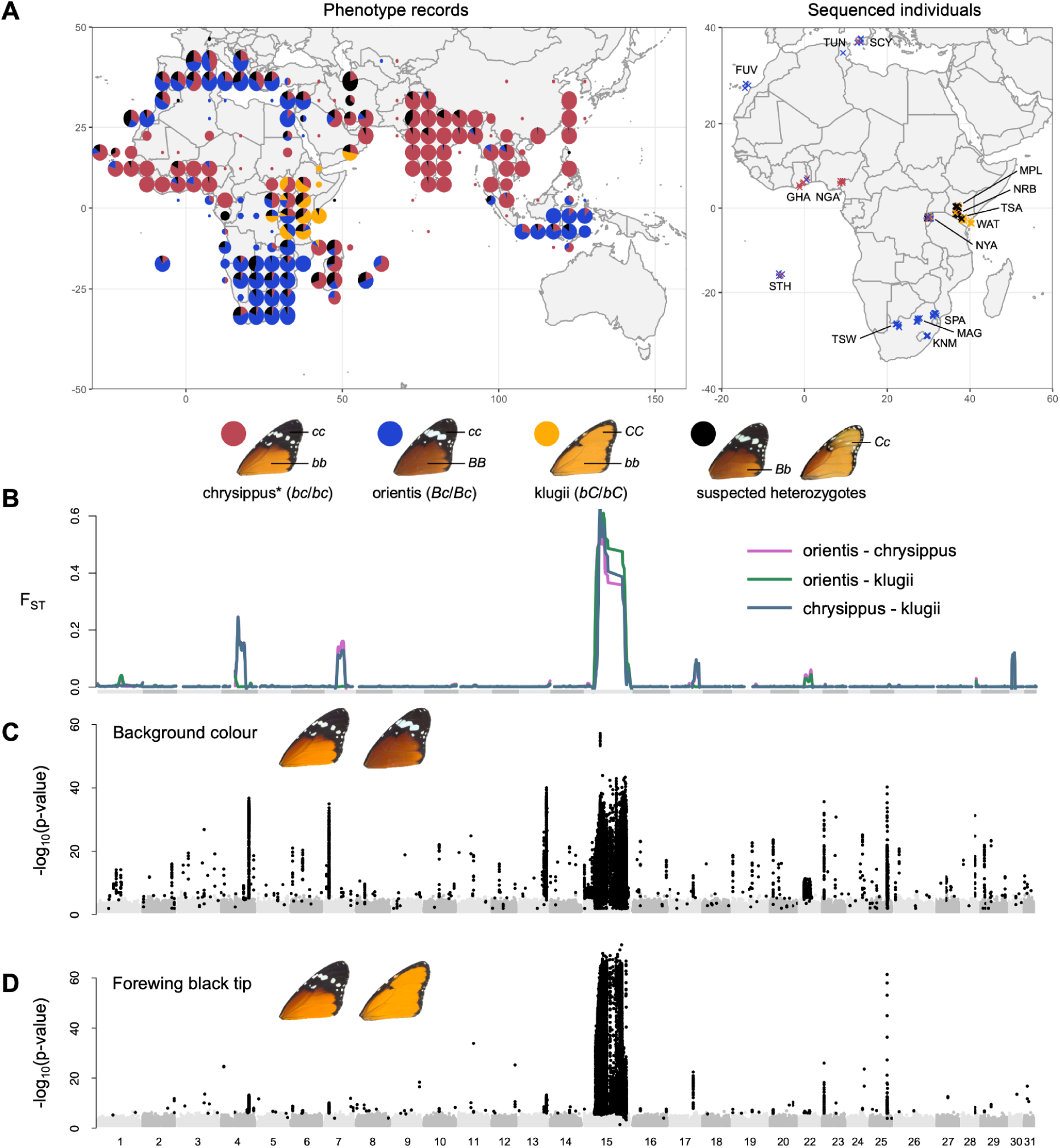
Broad geographic distribution of *Danaus chrysippus* morphs and our continent-wide sampling of individuals spans wing pattern variation that is underpinned largely by variation on chromosome 15. **A.** Geographic distribution of the three main *D. chrysippus* forewing morphs; chrysippus, orientis, and klugii as well as putative heterozygotes (data from manual curation of GBIF record and other scientific collections, following the approach in [44]; left) and sampled locations of *D. chrysippus* individuals used for genomic analysis in the present study (right). Public domain map from naturalearthdata.com. Morphs and their corresponding inferred genotypes at the B and C loci are shown below. Note that for our purposes the forewing morph ‘chrysippus’ includes the west-African morph ‘alcippus’, which has the same forewing phenotype but differs in its hindwing phenotype. **B.** Genome-wide patterns of *F*_ST_ between morphs (smoothed across discrete 50kb windows) highlighting significant differentiation along the length of the chr15 supergene region. **C-D.** Genome-wide associations between SNP variation and **C.** background coloration (individuals were scored as light, intermediate, or dark; N=172, following [44]) and **D.** forewing band presence/absence (individuals were scored as absent, partial, or present; N=172, following [44]) reflecting significant associations between this region and wing-pattern variation. Significant associations, as identified via permutation tests (p < 0.05), are represented by black points. The data underlying this Figure can be found in https://doi.org/10.5281/zenodo.14718778 and S2 Table.

Here we investigate haplotype diversity and evolution of the BC supergene using genomic data from 174 *D. chrysippus* individuals representing 14 regions across Africa and Southern Europe. We first confirmed that the supergene links loci controlling two distinct wing color pattern traits. We then describe haplotype diversity and identify additional divergent haplotype groups beyond those previously described. Comparable levels of genetic diversity in each haplotype group and across most of the discrete structural modules of the supergene argues against ongoing allelic turnover, and instead suggest a long-term polymorphism, probably driven by local adaptation. Perhaps most intriguingly, we find abundant evidence of recombination and gene flux between haplotype groups, with crossovers having occurred both between adjacent inversions and within individual inversions, leading to exchange of functional color pattern alleles. These results support a role for somewhat rare recombination in the diversification and long-term persistence of supergene haplotypes.

## Results

### Widespread sampling confirms that the BC supergene is associated with forewing color variation and is the main axis of genetic variation

We analysed resequenced genomes of 174 individual butterflies spanning the three core *D. chrysippus* forewing morphs, ‘chrysippus’ (note that for our purposes, this also includes form ‘alcippus’, which differs only in its hindwing), ‘orientis’, and ‘klugii’, as well as intermediate morphs (Fig. 1A; S2 Table). Our sampling includes areas of monomorphism for each morph: western Africa, South Africa and southeastern Kenya, respectively. We also sampled the hybrid zone (sampled in Rwanda and central Kenya, where all three forewing morphs and intermediates are found), and North Africa and the Mediterranean (where two of the three morphs, and intermediates are found). Twelve sequenced individuals were bred in captivity from parents of mixed or unknown origins and were not assigned to any geographic group. Pairwise *F*_ST_ between three monomorphic locations in South Africa (TSW population, orientis morph), Nigeria (NGA population, chrysippus morph) and Eastern Kenya (WAT population, klugii morph), is close to zero genome-wide, with the exception of a few large peaks, including the largest on chromosome 15 (chr15), as described previously (Fig. 1B; S2 Table)[45]. Genome-wide principal components analysis (PCA) for all autosomes excluding chr15 further supports minimal genetic structure outside of the BC supergene, except that samples from the small island of St. Helena, as well as captive-bred families each cluster separately from the remaining wild individuals (Figure A in S1 Text).

Genome-wide association (GWAS) analysis confirms that the BC supergene region on chr15 is associated with two forewing traits: pale/dark background wing color (also known as the B locus; Fig. 1C), and the presence/absence of the forewing black tip and white band (also known as the C locus; Fig. 1D). In addition to a large peak of association in the supergene, there are several peaks of association with pale/dark variation on other chromosomes, suggesting that other loci also contribute to this trait. A repeat of the GWAS using only samples from the hybrid zone (NYA, NRB, MPL populations, N=87) recapitulates the same general result (Figure B in S1 Text), confirming that the observed associations are not driven by correlated selection on other traits that follow a similar geographic distribution. The cluster of SNPs most strongly associated with background wing color fall between two genes: *yellow*, which is known to be involved in melanin synthesis and coloration in other insects [51], and the *achaete-scute complex protein T3-like*, which has been shown to be involved in wing scale development across Lepidoptera [52]. The SNPs most strongly associated with the presence or absence of the forewing black tip were found to fall within the copy-number-variable (CNV) region of the supergene (Figure C in S1 Text). These findings suggest that the B and C loci are distinct, and that the supergene is favoured because it maintains locally adapted combinations of alleles at these loci, and probably others among the ∼150 genes it encompasses, in the face of gene flow.

### Extensive genetic variation at the BC supergene

We previously described three divergent haplotype groups (sometimes called ‘alleles’) of the BC supergene, corresponding to the three forewing morphs, maintained by several inversions, intra-chromosomal translocations, and copy-number differences that suppress recombination [21,45]. We therefore expected that diploid genotypes across the BC supergene would form six distinct genetic clusters corresponding to the three homozygous and three heterozygous states. However, as we describe below, several lines of evidence suggest that there are additional divergent haplotypes, and possible recombinants. First, both a distance-based Neighbor-Net network and principal components analysis for the complete supergene (excluding the CNV region) show more complex genetic structuring than we expected (Fig. 2A, Figure A in S1 Text). Three clusters representing homozygous genotypes for the three previously-described haplotype groups are identifiable (Fig. 2A), and include individuals previously matched to these genotypes [21]. One large cluster of individuals is intermediate between the known chrysippus and klugii homozygotes (Fig. 2A) and includes known heterozygotes between chrysippus and klugii from crosses and suspected wild heterozygotes based on wing patterns. However, numerous other individuals are dispersed across the network at varying distances between the three homozygous clusters. We therefore hypothesised that in addition to heterozygotes among the three known haplotype groups, our sampling may include additional divergent haplotype groups and/or recombinants among the three common haplotypes.

**Figure 2.**
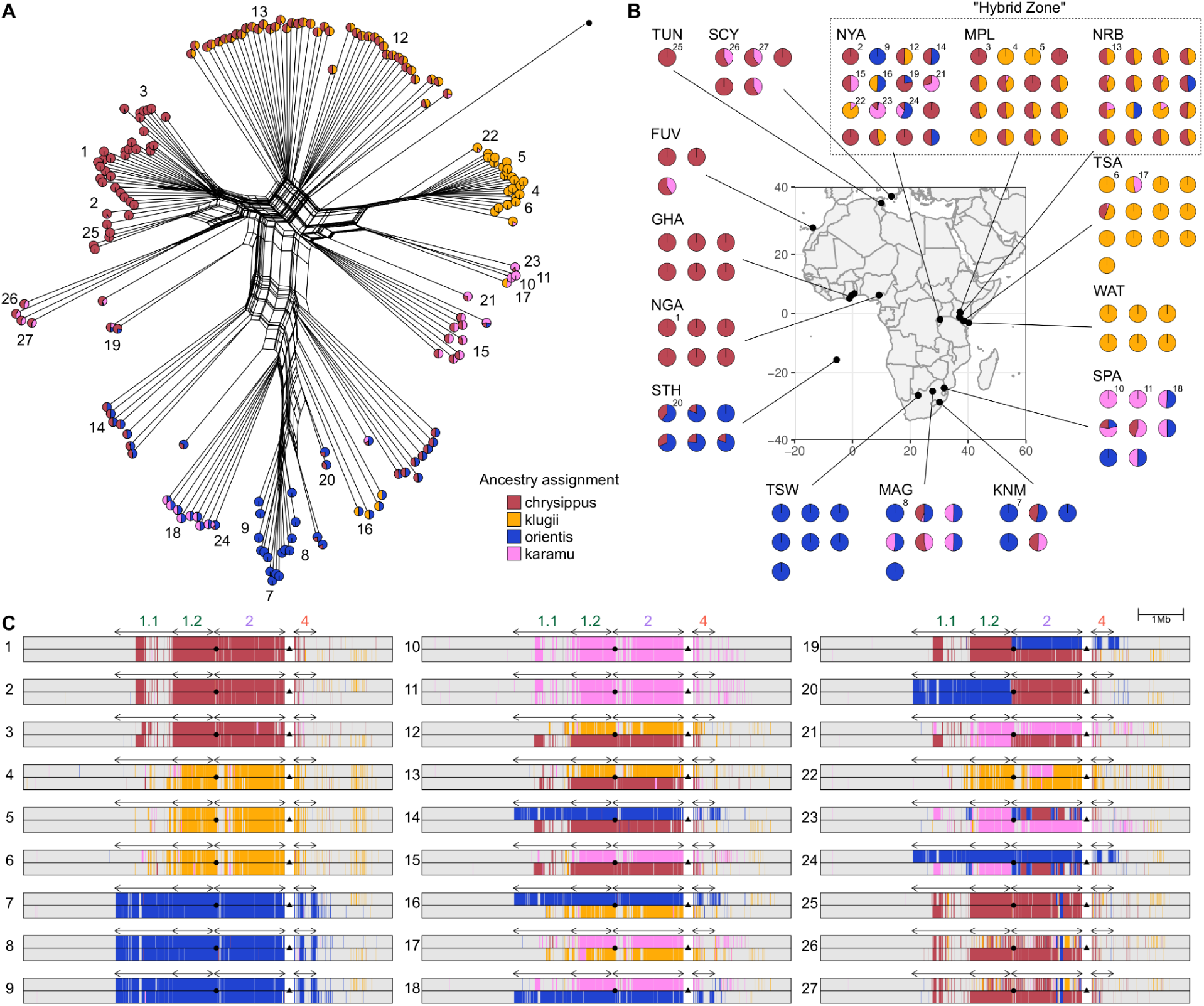
Genetic clustering and ancestry painting suggest additional divergent haplotype groups, and recombinants. **A.** Neighbor-Net network for unphased diploid genotypes across the BC supergene (excluding the CNV region). Each tip represents one diploid individual, and the network is constructed based on average pairwise genetic distances considering both haplotypes in each individual. We therefore expect ‘heterozygous’ individuals carrying two distinct haplotypes to be at intermediate positions in the network. Numbers indicate the 27 representative individuals for which ancestry painting is shown in Panel C. The single individual represented by a black dot is the *Danaus melanippus* outgroup. **B.** Localities for sequenced individuals (public domain map from naturalearthdata.com). Pie charts in panels A and B represent inferred ancestry components for each diploid individual from Admixture analysis with k=4 source populations (See Figures D-H in S1 Text for plots with other values of k). Note that for highly sampled localities, an arbitrary subset of sequenced individuals is shown in panel B. Numbered individuals correspond with those in Panels A and C. **C.** Ancestry painting across the central portion of chr15 including the BC supergene for 27 representative individuals, including homozygotes, heterozygotes and putative recombinants. See Figures I and J in S1 Text for ancestry painting for all individuals. The CNV region is excluded from the plot for convenience as it cannot be reliably genotyped (represented as a white gap). Arrows above the plots indicate the locations of inverted tracts 1.1 (1.3Mb). 1.2 (0.9 Mb), 2 (1.6 Mb) and 4 (0.5 Mb). The inferred location of the B locus (*yellow*) is indicated by a circle in the centre of each plot. The approximate location of the C locus is indicated by a triangle in the gap where the copy number variable region is found. The data underlying this Figure can be found in https://doi.org/10.5281/zenodo.14718778.

Given the complex relationships observed, we attempted to naively identify distinct haplotype groups using Admixture [53] analysis, in which each individual is modelled in terms of its ancestry components from *k* source populations. Because recombination should occur freely among haplotypes with the same structural arrangement, and should only be suppressed among those with different arrangements, the supergene region should comprise a set of semi-isolated sub-populations. We predicted that “heterozygous” individuals (i.e. carrying two divergent haplotypes) will appear as admixed, with approximately 50% ancestry contributions from two source populations, while homozygotes should be assigned 100% ancestry from a single source. All models with k=3 (matching our *a priori* hypothesis) and above captured the three previously-described haplotype groups corresponding to the klugii, chrysippus and orientis morphs. These are represented in clusters of homozygous (100% ancestry) individuals from eastern, western and southern Africa, respectively, while most hybrid-zone individuals appear heterozygous, with 50% ancestry proportions (Figures E-H in S1 Text). However, as described below, the models with *k*=4 to *k*=6, which all have lower cross-validation error than k=3 (Figure D in S1 Text), provide compelling evidence for the existence of additional divergent haplotype groups that had not been previously described.

We have chosen to present the ancestry assignments according to the model with *k*=4 sources in Fig. 2 (see pie charts in Fig. 2A and 2B), because our sampling included homozygotes for all four putative haplotype groups. The fourth source population is inferred to contribute ∼100% ancestry to two South African individuals, and ∼50% ancestry to several others. We therefore infer that a fourth arrangement of the supergene occurs in southeast Africa. This haplotype group is cryptic in the sense that it is associated with a forewing phenotype indistinguishable from that of orientis. Hereafter, we refer to this as the ‘karamu’ haplotype group.

In the model with *k*=5 sources (Figure G in S1 Text), the fifth cluster captures the previously described neo-W fusion - a lineage of chr15 haplotypes that arose recently in East Africa when a copy of chr15 from the chrysippus haplotype group fused to the female-limited W sex chromosome [45,49]. These females are each assigned ∼50% ancestry from this source, consistent with W being haploid in females. We previously showed that the neo-W-chr15 fusion occurred recently and spread to high frequency in southern Kenya over the past 2,200 years [45]. The high sequence similarity and complete lack of recombination of the neo-W explains how it is identifiable as a distinct genetic cluster despite its recent separation from the standard chrysippus haplotype group.

In the model with *k*=6 sources (Figure H in S1 Text), several individuals from North Africa and the Mediterranean are assigned to a sixth distinct cluster, suggesting yet another divergent haplotype group. We hypothesise that, similar to the karamu haplotype from southeast Africa, another distinct arrangement of chr15 likely occurs north of the Sahara. However, subsequent analyses (described below) suggest that none of our sequenced individuals are homozygous for this haplotype, limiting our ability to further investigate its status.

### Ancestry painting confirms additional haplotype groups, as well as recombinants

To investigate fine-scale haplotype ancestry and recombination across the BC supergene region, we applied two ancestry painting methods. These aim to assign ancestry for phased-inferred haplotypes based on pre-defined source populations, using either a hidden markov model [54] or defined windows of SNPs (see Methods for details). We used populations WAT, TSA and NGA as reference sets representing the klugii, orientis and chrysippus haplotype groups, due to their monomorphic clustering (Fig. 2B). In addition, we used the two individuals from the SPA population showing 100% karamu ancestry to represent this fourth haplotype group (Fig. 2B). Both ancestry painting methods agree strongly with our inferences from the Admixture analysis. All four haplotype groups occur as intact haplotypes in most individuals, with large numbers of both homozygotes (individuals 1-11 in Fig. 2C) and heterozygotes (individuals 12-18 in Fig. 2C). See Figures I and J in S1 Text for all individuals. Heterozygotes are particularly common in the hybrid-zone (populations NYA, MPL and NRB in Rwanda and central Kenya, Figures I and J in S1 Text).

Ancestry painting also identifies individuals with mosaic ancestry consistent with recombination between the divergent haplotypes. Most of these putative recombinant haplotypes appear to be simple chimeras that result from a single crossover located between two adjacent inversions (see for example individuals 19-21 in Fig. 2C). However, there are also instances of more fine-scale mosaic ancestry consistent with recombination within inversions (see for example individuals 22-25 in Fig. 2C). While gene conversion can lead to the transfer of small haplotype tracts within inversions (tract lengths vary across taxa but estimates suggest they are typically < 4kbp in length; [55]), this process cannot explain the scale of mosaic ancestry tracts we observe (tens to hundreds of kb). Instead, these patterns are more consistent with occasional double crossovers, which allow genetic exchange within the inversions without the formation of unbalanced gametes (but see Discussion). Some of these recombinant haplotype patterns appear multiple times in unrelated individuals, implying that they may be increasing in frequency in the population (see for example individuals 23-24 in Fig. 2C). Unsurprisingly, most recombinant haplotypes are found in the hybrid zone (Figures I and J in S1 Text). However, we were surprised to find two distinct recombinant haplotypes present in the genomes from the small island of St. Helena (STH population), where only one of the six sequenced individuals did not carry a recombinant haplotype (Figures I and J in S1 Text).

Ancestry painting failed to assign ancestry to one of the haplotypes in some individuals from North Africa and the Mediterranean (see for example individuals 26-27 in Fig. 2C). These are individuals that were assigned to the 6th cluster in the admixture analysis with k=6 sources (Figure H in S1 Text). This supports the existence of an additional divergent haplotype of the BC supergene that occurs north of the Sahara. Other members of this population carry a chrysippus-like haplotype, with some evidence for recombination with orientis (seen in individuals 25-27 in Fig. 2C). Because no individual is homozygous for the additional divergent haplotype, we were unable to include it as a source for ancestry painting, and it was not included in our subsequent analyses.

In summary, our findings suggest that there are (at least) six divergent haplotype groups of the BC supergene that experience suppressed recombination with respect to each other. Three were previously assembled and found to have distinct arrangements (chrysippus, klugii and orientis); one results from fusion of chr15 to the W chromosome (neo-W-chrysippus); and two are newly identified here based on genetic clustering, but probably also represent distinct structural arrangements (karamu and the unnamed haplotype from North Africa and the Mediterranean) (Table 1).

**Table 1.** Six divergent haplotype groups of the BC supergene.

| Haplotype Name | Geographic region(s) | Background color (B locus allele) | Forewing tip and band (C locus allele) | Conventional name of morph |
| --- | --- | --- | --- | --- |
| chrysippus | West Africa, North Africa & Mediterranean, Asia | Pale (*b*) | Present (*c*) | chrysippus, alcippus |
| klugii | East Africa | Pale (*b*) | Absent (*C*) | klugii |
| orientis | Southern Africa | Dark (*B*) | Present (*c*) | orientis |
| karamu | Southeast Africa | Dark (*B*) | Present (*c*) | orientis |
| neo-W-chrysippus | East Africa (Hybrid Zone) | Pale (*b*) | Present (*c*) | N/A (never homozygous) |
| Unnamed | North Africa & Mediterranean | Unknown (no homozygotes) | Unknown (no homozygotes) | Unknown (no homozygotes) |

### Diversity and divergence patterns are consistent with long-term polymorphism

One possible explanation for the coexistence of multiple haplotypes associated with the same wing phenotype in the same population (e.g. karamu and orientis haplotypes in the SPA population) is that we are witnessing an ongoing ‘allelic turnover’ event [42], in which a new haplotype is replacing an older one. If a new haplotype arose through a novel structural rearrangement, it is expected to have undergone a bottleneck of N=1 at its conception [3], which would result in dramatically reduced diversity immediately following its origin. Older haplotypes should have had the opportunity to recover diversity following their initial bottleneck, but are nevertheless still expected to have lower diversity than the genomic background level because inversions effectively divide the species into sub-populations with lower effective population size across the inverted region (even for the ancestral orientation) [3]. We therefore expected that all haplotype groups may show reduced within-group diversity within the inverted regions, but we hypothesised that a more pronounced reduction may be observed in those with derived inversion orientations: orientis and chrysippus [21] and possibly the newly identified karamu haplotype. Consistent with expectations, diversity is lower within the inverted regions of chr15 compared to collinear regions in all four haplotype groups (Fig. 3b). The extent of diversity reduction is notably similar in the four groups, with the exception that orientis has lower diversity in inverted region 1.1, which is known to be uniquely inverted in orientis. No haplotype group has diversity approaching zero, as would be expected under rapid, recent allelic turnover, so these patterns are more consistent with a long-term polymorphism at the BC supergene.

**Figure 3.**
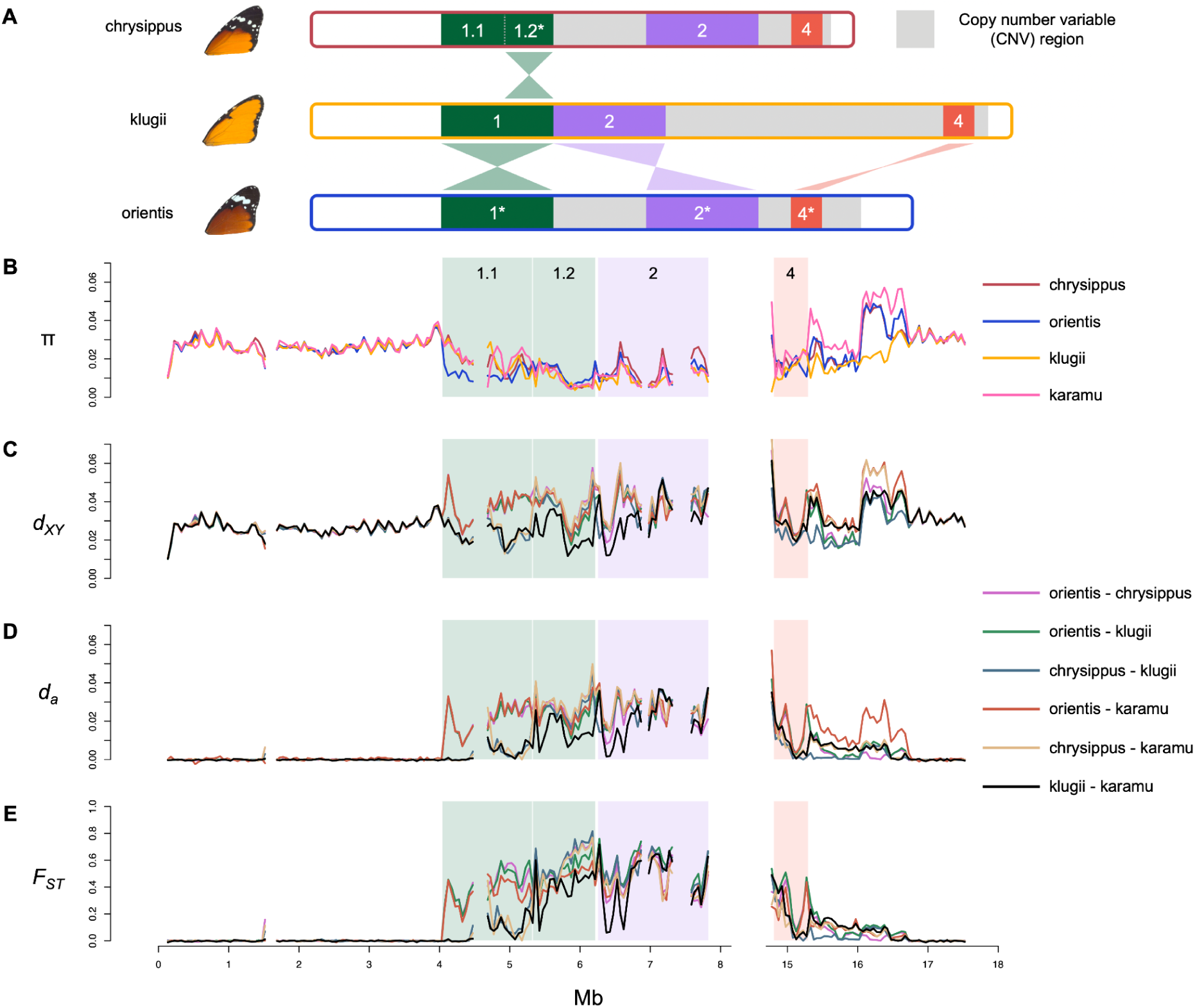
The supergene region has reduced genetic diversity in all haplotype groups and elevated genetic differentiation between them, but no evidence for a very recent origin of any haplotype. **A.** A graphical representation of the structural variation across the chrysippus, orientis, and klugii haplotypes described in [21]. The additional divergent haplotypes described in the present study have not yet been assembled with long reads, so their structures remain unknown. The three previously assembled haplotypes evolved through rearrangements of four main regions (indicated with different colors; regions with an inverted orientation relative to the ancestral state are indicated with *). Regions 1.1, 1.2, 2, and 4 remain in single-copy and can be reliably aligned and genotyped. **B-E.** Population summary statistics computed in non-overlapping 50kb windows for representative populations of chrysippus (NGA population), orientis (TSW), klugii (WAT) and karamu (two homozygous individuals from the SPA), plotted across chr15 (excluding the CNV region). Note that the reference genome is from a klugii haplotype, hence regions 1 and 2 are adjacent. Statistics plotted are: **B.** Nucleotide diversity (π) **C.** absolute pairwise divergence (*d_XY_*), **D.** net pairwise divergence (*d_a_*), and **E.** pairwise differentiation (*F_ST_*). The data underlying this Figure can be found in https://doi.org/10.5281/zenodo.14718778.

Genetic divergence and differentiation between haplotype groups are also consistent with relatively ancient origins. Elevated *F_ST_* and net divergence (*d_a_*) are restricted to the rearranged parts of the chromosome (Fig. 3), consistent with recombination suppression being restricted to the supergene (note that regions 1.1 and 4 are only inverted in orientis and are collinear in klugii and chrysippus). Similar levels of differentiation are seen between these three previously-described haplotype groups across most of the supergene (excluding region 1.1), which may indicate that the arrangements all occurred around the same time, although this pattern would also be expected under mutation-drift-flux equilibrium [3]. However, divergence and differentiation are both notably lower between klugii and the newly-described karamu haplotype group across most of the supergene, consistent with either a more recent divergence between these haplotypes, or a higher rate of gene flux between them. This is intriguing, because these haplotypes are associated with distinct wing phenotypes. This raises the possibility that gene flux between haplotype groups has allowed the exchange of functional alleles.

### Recombination between supergene haplotypes has exchanged wing color alleles

We set out to test whether the evolution of the karamu haplotype may have arisen through recombination between haplotype groups. Specifically, we applied topology weighting [56] using TWISST2 to ask whether there are tracts in which karamu is more closely related to orientis than to klugii. This revealed that karamu clusters broadly with klugii throughout the length of the supergene, but there are multiple narrow tracts at which it clusters more closely with orientis, particularly in inverted region 2 (Fig. 4A). Ancestry painting using Loter [54] confirms that one such narrow tract is a 20kb region containing the gene *yellow* and part of its promoter, including nine of the ten SNPs most strongly associated with wing background coloration according to our GWAS (Fig. 4B). This therefore suggests that incorporation of the orientis-like allele at *yellow* (i.e. the *B* allele at the B locus) through recombination causes the karamu haplotype to produce a dark wing color phenotype matching that of orientis, despite otherwise being more klugii-like in its overall ancestry. We hypothesise that a similar allelic exchange at the C locus causes karamu individuals to share the forewing black tip phenotype of orientis, but we are unable to test this hypothesis, as this trait maps to the CNV region in which genotyping and ancestry assignment are unreliable.

**Figure 4.**
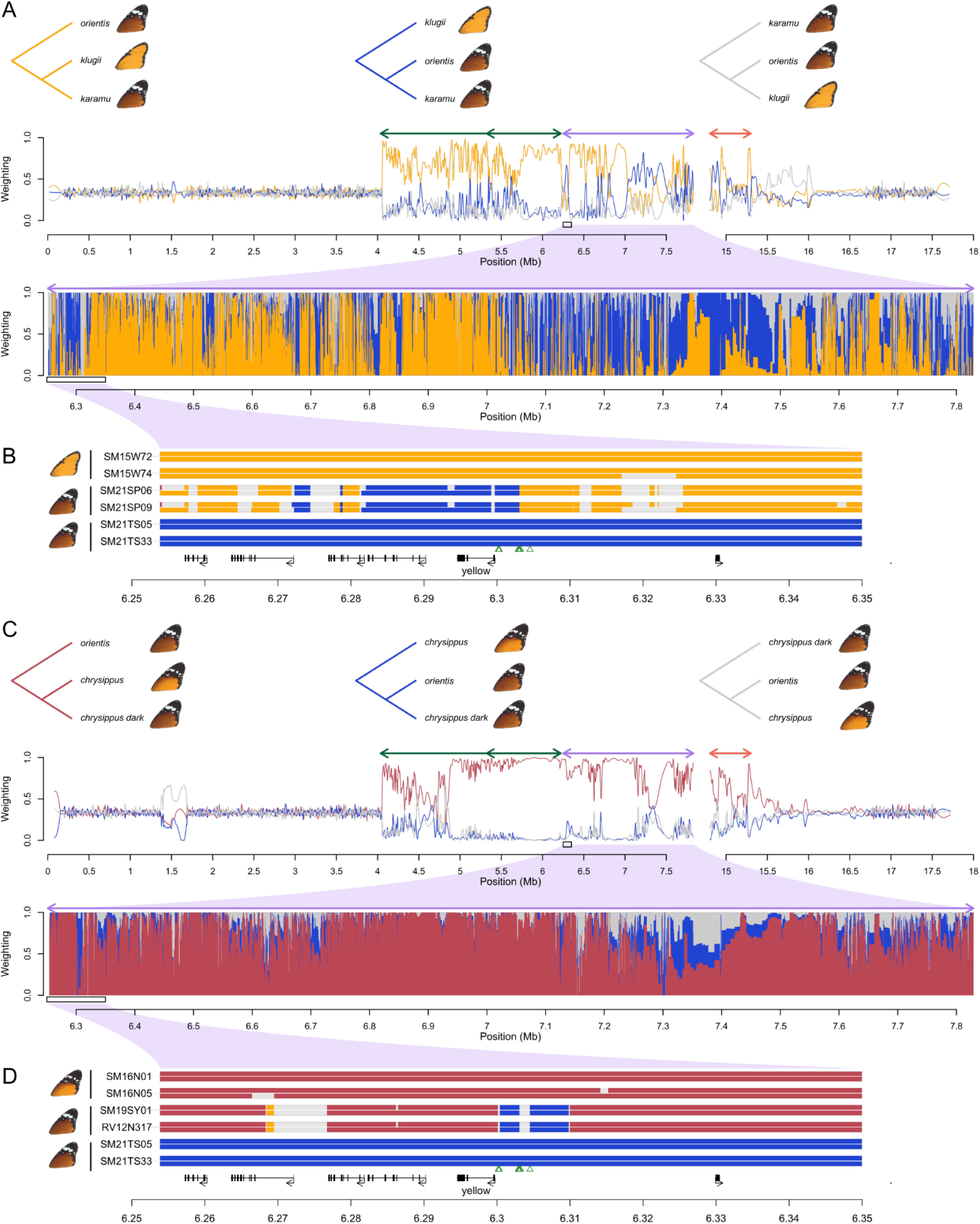
Two independent cases of recombination within the BC supergene with phenotypic consequences. **A.** Topology weightings across chr15 showing how the karamu haplotype is related to the klugii and orientis haplotypes. Upper panel shows three possible rooted genealogical topologies. Second panel shows weights for each topology along the chromosome, smoothed with a 20kb span. Arrows above the plot indicate the locations of inversions. Third panel shows unsmoothed topology weightings across a 1.5 Mb region corresponding to Inversion 2. **B.** Ancestry painting from Loter [54] across a 100 kb region within Inversion 2 showing ancestry tracts for two homozygous karamu individuals compared to two representative individuals homozygous for the orientis and klugii haplotypes. Coding regions are indicated below the plot, with the candidate gene for background coloration *yellow* indicated. Green triangles represent the top 10 SNPs for background color in our GWAS (Fig. 1C). There is evidence for recombination throughout the supergene region, and specifically in the vicinity of *yellow*, consistent with the hypothesis that orientis ancestry at this locus (i.e. the *B* allele) is associated with darker coloration in karamu individuals. **C** and **D.** A second example of recombination in the promoter of *yellow*. Plots are as described for panels A and B, except showing relationships between two individuals from North Africa & Mediterranean with a chrysippus-like haplotype (according to Admixture and ancestry painting), but dark background coloration. Again, orientis ancestry in the promoter region of *yellow* suggests that recombination allowed the transfer of the *B* allele into a different genetic background, causing darker wing coloration. The data underlying this Figure can be found in https://doi.org/10.5281/zenodo.14718778.

We identified a second probable case of recombination affecting background coloration. Several individuals from North Africa and the Mediterranean are homozygous for chrysippus-like haplotypes according to chromosome painting (see for example individual 25 in Fig. 2C), but have dark background coloration similar to that seen in orientis. We again tested for fine-scale mosaic ancestry in two of these individuals using topology weighting and ancestry painting. A very similar pattern to the karamu case is observed: these individuals cluster strongly with chrysippus throughout the supergene, but carry narrow orientis-like tracts (Fig. 4C), including a 10kb tract in the promoter region of *yellow* encompassing all ten SNPs most strongly associated with dark background coloration (Fig. 4D). This represents a distinct event in which orientis-like alleles (i.e. the *B* allele at the B locus) have been incorporated into a different genetic background through recombination within an inversion, leading to an altered phenotype.

### Historial gene flux also occurred between the major supergene haplotypes

Based on the extensive evidence for recombination observed, we hypothesised that even the three originally described haplotype groups (orientis, klugii and chrysippus) may have been historically shaped by gene flux. To explore this possibility, we first ran another topology weighting analysis to examine fine-scale relationships between these three groups. This generally agrees with the supergene structure: in each rearrangement, the predominant genealogy clusters groups according to whether they carry the rearranged or ancestral arrangement (Figure L in S1 Text). However, there is heterogeneity in these relationships, and occasionally, complete switches to a different relationship within each rearranged region (Figure L in S1 Text). These patterns are consistent with historical gene flux after the inversions occurred.

We next examined patterns of linkage disequilibrium (LD) between the three pairs of haplotypes. In each case, clear regions of high LD correspond with the previously described rearrangements that differentiate each pair [21] (Figure M in S1 Text). There is also quantitative variation in the extent of LD within inverted tracts, which could be indicative of variation in the extent of gene flux across the chromosomes, but may also reflect other factors such as variation in effective population size. Overall, the generally high LD within inversions, along with high absolute genetic divergence across the length of the supergene (Fig. 3), imply that any recent gene flux between these haplotypes has occurred at a low rate.

Finally, we attempted to quantify gene flux between the three major haplotype groups by fitting isolation-with-migration (IM) models, which are conventionally applied to model species divergence with gene flow. Here haplotype groups are modelled as panmictic populations that became isolated at some point in the past, and gene flux between the distinct haplotype groups is modelled as ‘migration’ [36]. IM models were fitted using gIMble [57], with separate models for each rearranged region and each pair of haplotype groups, because not all regions are inverted between all pairs. For example, regions 1.1 and 4 are inverted in orientis only, so we expect approximately free recombination between klugii and chrysippus in these regions. Of the eight pairwise tests in which the regions are inverted, three had best-fitting models without gene flux, while the remaining five had estimated effective migration rates (*m_e_*) ranging from 2.42x10^-11^ to 3.09x10^-6^ (this equates to a range of 3.02x10^-5^ to 4.89 effective migrants per generation (*M_e_*) assuming *M_e_ = m_e_4N_e (recipient)_*; Table 2; Figure N in S1 Text). Higher rates are generally seen between klugii and orientis, but there is no apparent trend with the size of the rearranged region. Of the four tests that involved collinear regions, two had the highest estimated effective migration rates, and two had low or no inferred gene flux. This is probably because these latter two collinear tracts (specifically region 1.2 between chrysippus and orientis, and region 2 between klugii and chrysippus) are nevertheless physically translocated with respect to each other (Fig. 3A), probably impairing correct meiotic pairing. Taking these results together, we conclude that there is considerable evidence for history of gene flux in at least some of the rearranged regions of the supergene, but our estimated rates of gene flux should be interpreted with caution given the small size of the regions and resulting sampling noise and reduced inference power (we note that one model failed to optimise), and the modelling restriction of unidirectional gene flux [57].

**Table 2.**
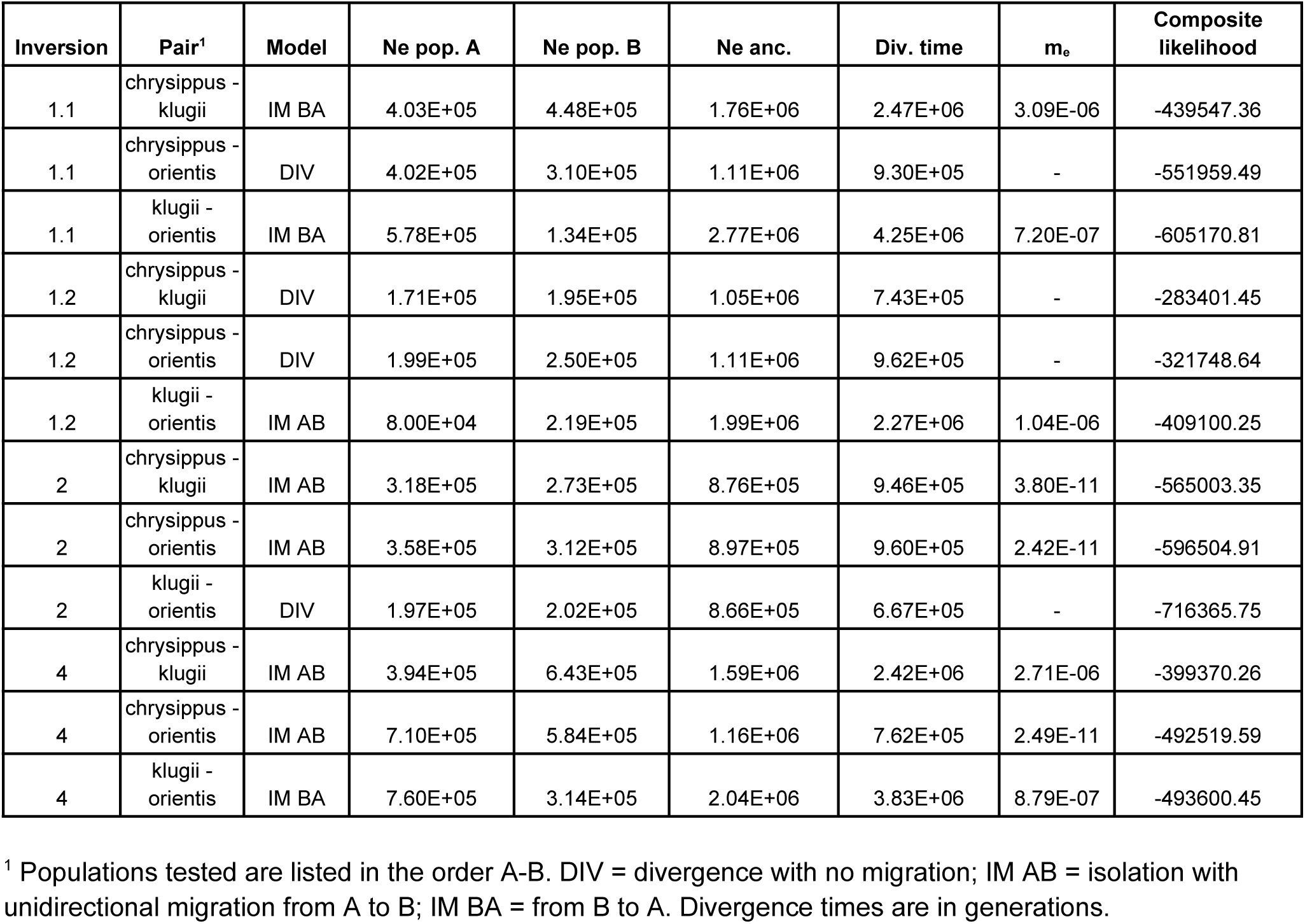
Summary of gIMble results highlighting the best fitting models for each region and haplotype pair combination.

## Discussion

Structural variants that suppress recombination have been found to contribute to local adaptation and the maintenance of complex phenotypic polymorphisms. However, to understand how these variants persist and change over time, it is necessary to go beyond simply associating genotype and phenotype, and to investigate how sequences evolve within these genomic regions. Here we show that the BC supergene of *Danaus chrysippus*, which underpins wing color pattern variation, displays surprising haplotype diversity, with a greater number of divergent haplotype groups than there are described phenotypic morphs. Counterintuitively, this diversity is partly fuelled by recombination. Our findings suggest that supergene evolution is a highly dynamic process, raising a number of further questions.

A paradox of our study is that we report evidence for both historical and recent recombination in a genomic region that was first identified because of its role in suppressing crossing over. In reality, the realised rate of gene flux through recombination in the BC supergene must be low, otherwise we would not have been able to describe the highly diverged haplotype groups, nor to identify recombinants between them. Indeed, although a considerable fraction of our sequenced individuals carry haplotypes showing evidence for recent recombination, many of the recombination breakpoints are shared among individuals, implying that the observed recombinant haplotypes stem from a smaller number of recombination events. Moreover, several recombination points are between adjacent inversions, leaving the inversion haplotypes intact. The modular structure of the supergene means that single crossovers between adjacent inversions can generate modular ancestry mosaics without the production of unbalanced gametes. However, we also observe more complex mosaic ancestries within inversions in some individuals, as well as evidence for long-term gene flux, consistent with some recombination occurring within inversions. A large proportion of this may be attributable to gene conversion, which has been shown to be unimpeded within inversions in *Drosophila* [28–32,38,40]. However, gene conversion cannot easily explain the exchange of long tracts that we observe, such as the 10-20kb tracts around the *yellow* locus. It is equally difficult to explain these by double crossover events, which should be impeded due to crossover interference in inversions of 1-2Mb in length. Indeed, chromosome map lengths in the butterflies *Heliconius melpomene* and *Vanessa cardui* average around 50 cM [58,59], implying an average of one crossover per meiosis per bivalent, and in *Leptidea sinapis* only chromosomes >20Mb (longer than any chromosome in *D. chrysippus*) have map lengths consistent with frequent double crossovers [60]. An alternative explanation for the large tracts of recombinant ancestry within inversions is that these may have been facilitated by recurrent inversion or “toggling” of inversion orientations [61]. Toggling may be common in inversions flanked by segmental duplications due to recurrent non-allelic homologous recombination (NAHR) events [61], and could serve as a bridge for recombination between inverted haplotypes. Another line of evidence consistent with the toggling hypothesis is a lack of the characteristic “suspension bridge” pattern of divergence reflecting greater recombination suppression near the breakpoints, as seen in some other inversions [35]. Importantly, once a large tract of ancestry has been exchanged (either via double crossover or inversion toggling), the recombined tract is no longer inverted with respect to the haplotype it has invaded. Hence, more complex fine-scale ancestry mosaics such as those we describe can emerge through unrestricted single crossovers in subsequent generations.

Our findings do not provide direct evidence of the potential costs or benefits of recombination within the supergene region. The simplest models of supergene evolution imply that recombinant haplotypes have reduced fitness due to broken associations between co-adapted or locally adapted alleles. It is notable that one particular combination of color pattern alleles is absent in our data set: haplotypes carrying the *B* (dark) and *C* (absence of forewing tip) alleles. All individuals inferred to be homozygous *BB* based on the best GWAS hit for the B locus (position 6,300,183 on chr15) have a full forewing black tip (implying a *cc* genotype at the C locus). The rarity of the *BC* haplotype in the wild despite its occasional production in breeding experiments has been noted previously [62], and is suggestive of selection against this particular allelic combination. Another intriguing observation is that the population with the highest rate of recent recombinants is the small island of St. Helena, where five of six individuals carry one of two different recombinant haplotypes between chrysippus and orientis. We hypothesise that this population is the result of a recent colonisation bottleneck and that the abundance of recombinant haplotypes may result from relaxed purifying selection. On the other hand, it is also likely that certain recombinant haplotypes are occasionally more fit than the prevailing haplotypes, either because they provide new allelic combinations (including at loci other than B and C) associated with a fit phenotype [5,22,37], or because they allow purging of deleterious mutations [23]. Our finding that several haplotypes with the same complex ancestry mosaics appear in multiple individuals may imply a process of allelic turnover, in which an existing ‘allele’ (or haplotype group) is in the process of being replaced by a fitter recombinant one. Indeed, the complex pattern of relationships among even the established haplotype groups suggests that recombination and allelic turnover may have occurred repeatedly throughout the evolutionary history of the BC supergene. As a result, it is likely that none of the established haplotype groups we describe here closely resemble the ‘original’ haplotypes captured when the rearrangements first occurred. A process of dynamic turnover has been suggested in a different supergene system in *Papilio* butterflies [42], and may prove with further work to be the norm for supergenes, as it is for sex chromosomes in some taxa [63].

A puzzling feature of the BC supergene is the apparent long-term persistence of multiple haplotype groups with (apparently) the same phenotypic effects. Specifically, the newly described karamu haplotype found in eastern parts of South Africa is associated with a color pattern indistinguishable from that of the orientis haplotype found throughout South Africa. Although we initially hypothesised that this may represent a case of ongoing allelic turnover (in which the karamu haplotype may eventually replace the orientis haplotype), this is not supported by the similarly high level of diversity seen in both haplotype groups, implying that neither has experienced a recent or ongoing selective sweep. An alternative explanation is that both of these haplotypes may be adapted to different geographic regions by influencing traits other than color pattern. It is certainly possible that the BC supergene region, which contains approximately 150 protein-coding genes, contributes to other important ecological traits. Simulations have shown that such adaptations may continue to accumulate after inversions have become established [11]. It is notable that the karamu haplotype appears to have arisen through recombination between the southern orientis and eastern klugii haplotypes, and appears to be most common in a geographic region intermediate between these two (though our sampling is too sparse to be sure). It is therefore plausible that this haplotype confers an adaptive advantage by combining an optimal warning pattern for southern Africa with other environmental adaptations suited to east Africa. A similar scenario may explain the existence of a dark-colored morph in North Africa and the Mediterranean with a haplotype that is highly similar to that of the pale-colored chrysippus morph, but carries orientis-like variants at the promoter of *yellow* that result in dark coloration. In this case, it may be that recombination allowed for allelic replacement at one locus while maintaining the broader locally-adapted haplotype. Further investigation into the ecology and fitness of different genotypes and phenotypes is required to test these hypotheses.

Another factor that may shape haplotype diversity is vulnerability of inversions to the accumulation of deleterious mutations. Both theoretical and empirical studies have shown that inversions can accumulate excess deleterious load as a result of their lower effective population size compared to the rest of the genome and their reduced opportunities for recombination [3,23,24]. This may drive turnover if new or recombinant haplotypes have lower mutational load. It has also been suggested that load may promote polymorphism through associative overdominance, in which heterokaryotypes carrying different sets of deleterious recessive alleles have increased fitness relative to homokaryotypes due to masking of the deleterious recessives [23]. While examples of heterokaryotype advantage are known [24], it is difficult to prove that associative overdominance is the cause, especially given that the conditions under which it is likely to evolve are highly restrictive [25]. In *D. chrysippus*, there is no compelling evidence for heterozygote advantage: the three most common haplotype groups each occur in large regions of monomorphism, and broad clines suggest that polymorphism in the hybrid zone reflects a balance between selection (local adaptation) and extensive dispersal [44] rather than heterokaryotype advantage. Further investigation is needed to specifically rule out heterokaryotype advantage, especially for the newly identified karamu haplotype and the other divergent haplotype found north of the Sahara, which are only known from polymorphic locations. It is worth noting that the recent fusion of chr15 (chrysippus haplotype) to the W chromosome in East Africa [45] represents the origin of yet another divergent haplotype that is restricted to the heterozygous state (butterfly females are ZW and do not undergo recombination), and therefore would be sheltered from the effects of deleterious recessives.

The additional divergent haplotype groups described in the present study were identified using only short-read sequencing data, so we cannot confirm whether each of these are also maintained by unique rearrangements of chr15, though this seems highly likely. Using short reads aligned to a single reference also means that insertion, deletion and copy number variants were not considered, which probably explains our inability to narrow down the C locus. Future work using genome graph approaches will provide an improved understanding of the relationships between structural variation, haplotype diversity and phenotypic variation.

We hypothesise that the unusual physical structure of *Danaus* chromosome 15 has contributed to the emergence of divergent haplotypes. The chromosome carries a large copy-number-variable region which began to grow around 7.5 MYA, and now comprises over a third of the chromosome in the klugii haplotype [21]. This region, which is also rich in transposable elements, creates fertile ground for the emergence of new divergent haplotypes through two mechanisms of recombination suppression [21]: It could directly prevent crossing over [64] or indirectly promote recombination suppression by propagating new inversions through non-allelic homologous recombination (NAHR). Whether the physical peculiarities of this chromosome also facilitated its recent fusion to the W chromosome in East Africa remains to be determined. It has been suggested that the occurrence of holocentric chromosomes (centromeres are dispersed across the length of the chromosome) and inverted meiosis (sister chromatids separate before homologous chromosomes) in the lepidoptera might allow for more rapid evolution of chromosomal rearrangements on average in this insect order [65]. However, a recent analysis of chromosome-scale genomes from 210 lepidopterans revealed that both inter- and intrachromosomal rearrangements are rare overall outside of a few highly rearranged lineages (not including *Danaus*) [66]. The tendency for *Danaus* chromosome 15 to experience rearrangements is therefore more likely a feature of the individual chromosome than of the lineage. This echoes a similar pattern seen in the avian chromosome 4, which has undergone both ancient rearrangements (possibly associated with an ancient supergene [67]), and more recent fusions and further rearrangement in the formation of neo-sex chromosomes [68–70]. It is notable that the *D. Chrysippus* BC supergene, the *Heliconius numata* P supergene [24] and the fire ant social supergene [6] all share the feature of multiple adjacent non-overlapping inversions that appear to share breakpoints. This modular organisation means that crossovers that occur between two inversions are not suppressed, allowing the distinct modules of the supergene to evolve somewhat independently. Modular arrangements may simply be a side-effect of ‘reuse’ of inversion breakpoints (e.g. in repeat-rich regions), but it is plausible that supergenes with this structure may be more likely to persist long-term if it avoids some of the costs of complete recombination suppression.

In conclusion, we have shown that a supergene with an apparently simple phenotypic association shows unexpected diversity at the haplotype level. Our findings bring to light the nuanced relationship between supergenes and recombination, in which incomplete recombination suppression can fuel haplotype diversification, and possibly support long-term persistence. Further work is needed to understand the selective and ecological drivers underlying the observed diversity. Our study adds to a growing body of work revealing the indirect effects of structural variation on the evolution of surrounding sequences [24,36,71,72]. Further work across a diverse range of taxa, along with population genetic modelling will help to uncover the generality of these phenomena.

## Methods

### Data collection, whole genome sequencing and genotyping

We produced new whole genome sequence data (Illumina, 150 bp paired-end) from 130 butterflies and combined this with previously sequenced data from additional 44 individuals [21,45,73], resulting in data from 162 wild-caught *Danaus chrysippus* butterflies representing each of the main color morphs across Africa and the hybrid zone (Fig. 1A, S2 Table) and 12 captively reared individuals. In addition, publicly available sequencing data from one *Danaus melanippus* individual was used in analyses that required an outgroup. Sample collection was conducted with the permission of landowners and under appropriate permits where relevant: Wildlife division of Ghana Forestry Commission: 0174076/024992; National Commission for Science and Technology, Kenya: NACOSTI/P15/3290/3607, NACOSTI/P15/2403/3602; Ministry of Education, Rwanda: MINEDUC/S&T/459/2017; Mpumalanga Parks and Tourism Agency, South Africa: MPB5667; Northern Cape Department of Environment and Nature Conservation, South Africa: FAUNA0615202; Environment & Natural Resources Directorate, St. Helena Government: EMDEP006/17; Council of Fuerteventura, Spain: Permit 604 (2017). All sample information, including accession numbers, are provided in S2 Table.

For samples newly sequenced in this study, the protocol followed our previous work [21]. Briefly, genomic DNA was extracted from ethanol-preserved tissue using the Qiagen DNEasy Blood and Tissue Kit (Qiagen, Redwood City, CA). Illumina library preparation and sequencing (150bp, paired-end) was performed by Novogene (Cambridge, UK), using a Novaseq 6000 instrument.

Reads were mapped to the Dchry2.2 reference genome (ENA accession GCA_916720795.1; [73]) using BWA mem v0.7.17 [74] with default parameters, and PCR duplicates were removed using PicardTools v2.11.1 (https://broadinstitute.github.io/picard/). Indel realignment was then carried out using GATK v.3.8 [75]. To identify any substantial variation in read depth across the dataset, mean read depth was calculated using Mosdepth [76]. One individual was found to have substantially higher read depth (SM18W01) and as a result the bam file for this individual was randomly downsampled to 30% of its original size using SAMtools ([77]; samtools view -s 0.30) to eliminate biases in genotyping. Genotyping was carried out using BCFtools [78], with genotypes on the Z chromosome (chr1) called by defining ploidy based on known sex (i.e. females as haploid and males as diploid). Genotypes were then filtered using BCFtools to remove any positions called fewer than twice and to leave only genotypes with an individual depth of >= 7 and with genotype quality of >=30. Additional filtering removed SNPs with >10% missing data (corresponding to a haploid count of 313 across autosomes and 224 on the Z chromosome), was performed for most remaining analyses, except when otherwise specified.

### Population structure, diversity and divergence

Genomic variation across our dataset, spanning different regions of the genome, was assessed by producing PCAs from different sets of SNPs using PLINK [79]. Two SNP sets were used, representing (i) the autosomal regions of the genome excluding chr15 as well as the Z sex chromosome chr1 (14,548,985 SNPs), and (ii) rearranged regions 1, 2, and 4 of the BC supergene on chr15, described in [21] (173,998 SNPs). Genome-wide nucleotide diversity (π), *F_ST_*, and *d_XY_*were computed in non-overlapping 50kb windows across the genome using the script *popgenWindows.py* (github.com/simonhmartin/genomics_general release 0.2) and net divergence (d_a_) was calculated from this output (calculated as 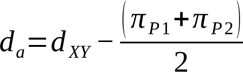 where P1 and P2 represent Population 1 and Population 2, respectively), following equations 10.5 for *π*, 10.20 for *d_XY_*, and 10.21 for *d_a_* in Nei et al. [80]. For each pair of haplotypes compared, only sites that had non-missing genotypes in both haplotypes were considered when computing the proportion of differences.

### Genome-wide association

The association between SNP variation and phenotypic variation was analysed using a genome-wide association analysis implemented in PLINK across all chromosomes. Phenotypes of scanned wings from 172 individuals were scored following [44] based on whether the background color was pale (1), intermediate (2), or dark (3), and whether the forewing band was absent (1), partial (2), or present (3) (wings were not available to phenotype for two individuals resulting in their omission from the GWAS). Empirical p-values were computed using the default adaptive permutation method implemented in PLINK. To confirm that geographic sampling structure alone does not underpin the patterns of association observed across the genome we re-ran the GWAS using only the 87 individuals from the hybrid zone for which we had phenotypes (populations NYA, NRB, and MPL).

### Visualising variation in the BC supergene using Neighbor-Net

To visualise genetic relationships and clustering across the BC supergene, we generated a phylogenetic network based on pairwise genetic distances among diploid individuals using the Neighbor-Net algorithm [81], implemented in SplitsTree [82]. Pairwise distances were computed using the script distMat.py (github.com/simonhmartin/genomics_general release 0.2), which computes the average pairwise distance between the four pairs of sequence haplotypes for each pair of diploid individuals. This analysis was computed using modules 1, 2 and 4 of the supergene region [21]. Input genotypes were filtered using the script filterGenotypes.py (github.com/simonhmartin/genomics_general release 0.2) to retain sites where at least 50% of individuals had non-missing data (after depth and quality filtering described above), where fewer than 75% of individuals were heterozygous, and where the minor allele was present at least twice.

### Admixture

We aimed to naively determine the number of distinct haplotype groups of the supergene by modelling each individual in terms of its ancestry from a fixed number of source populations. To this end, we ran Admixture v1.3.0 [53] on a concatenated dataset of loci from across the supergene modules 1, 2, and 4 (i.e. excluding the CNV region). Admixture was run specifying a range of clusters from k=3 to k=12 (Figure D in S1 Text) and 20-fold cross-validation error (--cv=20) was computed.

### Phasing

Phasing was carried out to assist with ancestry painting using two consecutive tools, Whatshap v1.1 [83], a read-based phasing software, and Shapeit4 v4.2.2 [84] a statistical phasing software using default parameters.

### Ancestry painting and additional phase correction

To visualise genetic clustering of phased haplotypes, we used two approaches for ancestry painting along the chromosome. Both approaches require defined reference individuals, which were selected based on the Admixture analysis above. Reference individuals used for each analysis are described in the Results.

The first approach was Loter [54], which uses a hidden Markov model to assign ancestry and identify breakpoints along the genome. First, phased genotypes were filtered using filterGenotypes.py (github.com/simonhmartin/genomics_general release 0.2) to remove any sites at which more than 60% of individuals are heterozygous, as such sites are likely to be caused by mis-mapping and could be prone to break tracts of ancestry. Haploid genotypes were output as counts of the minor allele (0 or 1), and Loter was run using the python API in a wrapper script (loter_wrapper.py), with options rate_vote=0, nb_bagging=20. Adjacent SNPs with identical ancestry scores were concatenated into tracts to reduce the output file size.

A second approach for ancestry painting that we call ‘distPaint’ was applied to confirm the Loter results, while providing a coarser visualisation at the whole-chromosome scale. This approach uses a sliding-window along the genome and assigns ancestry for each haplotype based on genetic distance (*d_XY_*) from the reference sets of individuals. A window size of 200 SNPs was used. Ancestry was assigned if this difference was significantly lower for one reference population according to a Wilcoxon Rank Sum test (p<=0.01), otherwise ancestry was set to undefined. This was implemented using a custom python script distPaint.py (github.com/simonhmartin/genomics_general).

Initial visualisation of the ancestry painting from both approaches revealed obvious cases of phasing errors (Figure K in S1 Text). Such errors could lead to false-positive inference of recombination events. For example, say an individual is phased into two haplotypes, that are then painted with ancestries “c-c-c-c-k” and “k-k-k-k-c” (where c and k represent tracts of chromosome assigned to reference populations chrysippus and klugii, respectively). This could either represent a case of an individual carrying two recombinant haplotypes in which both recombination events happened to occur at the same genomic region, or it could be a case of a single phasing error in an individual that in fact caries two common haplotypes (c-c-c-c-c and k-k-k-k-k). The latter is of course far more likely. We therefore applied a final heuristic phase correction approach to the inferred ancestries which attempts to maximise similarity among haplotypes at the whole-chromosome scale, thereby minimising inferred recombination events (Figure K in S1 Text). For each diploid individual, this heuristic approach iterates over each ancestry-painted tract in the chromosome (e.g. from Loter) and switches the phase if this would cause an increase in similarity of the two resulting haplotypes to one or more other haplotypes in the full dataset (considering all ancestry blocks up to and including the focal one). Because we are interested in identifying common shared haplotypes, we consider the average similarity of the top three most similar haplotypes in the rest of the dataset. The process is performed left-to-right and then right-to-left for each individual, and then repeated again starting from the first individual, up to a total of 20 iterations for Loter outputs and 500 iterations for distPaint.py outputs (the latter have fewer intervals, allowing for more rapid computation). This algorithm is implemented in the script phasepaint.py (https://github.com/simonhmartin/phasepaint).

### Topology weighting

To complement the ancestry painting described above, we analysed changes in relationships among supergene haplotypes across the chromosome (as an indicator of possible recombination), using topology weighting [56], implemented in TWISST2 (https://github.com/simonhmartin/twisst2). This approach infers genealogies and where they change along the genome, and outputs a ‘weighting’ for each possible topology for the relationships between defined groups of individuals in each genomic region. Input genotypes were filtered using the script filterGenotypes.py (github.com/simonhmartin/genomics_general release 0.2) to retain sites where at least 50% of individuals had non-missing data (after depth and quality filtering described above), where fewer than 75% of individuals were heterozygous, and where the minor allele was present at least twice. An outgroup was needed for polarisation, for which we used a single *Danaus melanippus* individual (S2 Table), following the alignment and genotyping procedures described above.

### Linkage disequilibrium analysis

Linkage disequilibrium (LD) between pairs of sites across chromosome 15 was quantified using the *r^2^* metric, computed using PLINK [79] with the flag --r2. The computation was run first on all 174 individuals, and then on three subsets representing each pair of reference individuals for chrysippus (population NGA), klugii (WAT) and orientis (TSW). SNPs with a minor allele frequency < 0.2 were excluded, as were all SNPs within the copy number variable region. Finally, SNPs were thinned randomly to retain ∼20,000 for each comparison. LD heatmaps were plotted using the LDheatmap package [85] in R.

### Modelling gene flux using gIMble

To detect and quantify gene flux between supergene haplotypes, we fitted isolation-with-migration models for individuals homozygous for the chrysippus, klugii, orientis alleles using the gIMble framework [57]. Under the IM model, gene-flux between supergene haplotypes is modelled as migration (gene flow) between panmictic populations. Six klugii individuals (WAT population), seven orientis individuals (TSW population), and six chrysippus (NGA population) were included in the analysis (S2 Table). Since gIMble carries out pairwise tests this resulted in six pairwise tests for each of the 4 regions tested. A VCF containing only data from these 19 individuals was first preprocessed using *gimble preprocess*. Next, intergenic bed files were produced by using bedtools intersect and bedtools subtract to extract and then exclude genic regions using the reference annotation file. *gimble parse* was then run for each combination of supergene region (i.e. regions 1.1, 1.2, 2, and 4, Fig. 3A) and pairwise comparison, followed by *gimble blocks*, *gimble windows*, *gimble info*, and *gimble tally*. *gimble optimize* was then run for each combination of region and pairwise test specifying each of three models, DIV - a divergence only model, IM_AB an isolation with migration model where gene flow occurs from population A into B, and IM_BA an isolation with migration model where gene flow occurs from population B into A, resulting in a total of 36 models. *gimble query* was then run to extract model summary information, including composite likelihood, for each model allowing us to determine the best fitting of the three models for each of the 12 region and pair combinations (full results are reported in S3 Table. For detailed commands and model parameters see archived GitHub repository: https://doi.org/10.5281/zenodo.14726309.

## Data availability

All raw sequencing reads are available via the European Nucleotide Archive (ENA) - Sample accession numbers are provided in S2 Table. Data underlying all figures, along with scripts to reproduce them are archived on the Zenodo Repository: https://doi.org/10.5281/zenodo.14718778. Commands and scripts for all data analyses are provided in an archived GitHub repository: https://doi.org/10.5281/zenodo.14726309.

## Acknowledgements

We thank the following people for assistance with sampling, breeding and acquisition of permissions: Rwanda - Constantin Sibomana and Jody Garbe; Kenya - Piera Ireri, Ivy Ng’Iru, Godfrey Etelej, David Smith, Anna Orteu, Jenny York, Owen McMillan, Valarie McMillan; South Africa - Jeremy Dobson; Ghana - Oskar Brattstrom; Spain (Fuerteventura) - Yeray Monasterio, David Smith; Tunisia - Roger Vila; Italy: Richard ffrench-Constant; St. Helena: David Pryce. We also thank Brian Charlesworth, Deborah Charlesworth, David Smith, Richard ffrench-Constant, Chay Graham, Thomas Decroly, Alexander Mackintosh, Domink Laetsch and Konrad Lohse for valuable input on this work.

## Supporting Information file legends

**S1 Text.** Supplementary Figures A-N

**S2 Table**. Sample information for all 174 *D. chrysippus* individuals and one outgroup *D. melanippus*, including sampling location, sex, phenotypes, and read accessions.

**S3 Table**. Full results of gIMble analyses.

