## Supplementary material for "Incomplete recombination suppression fuels extensive haplotype diversity in a butterfly color pattern supergene": S1 Text

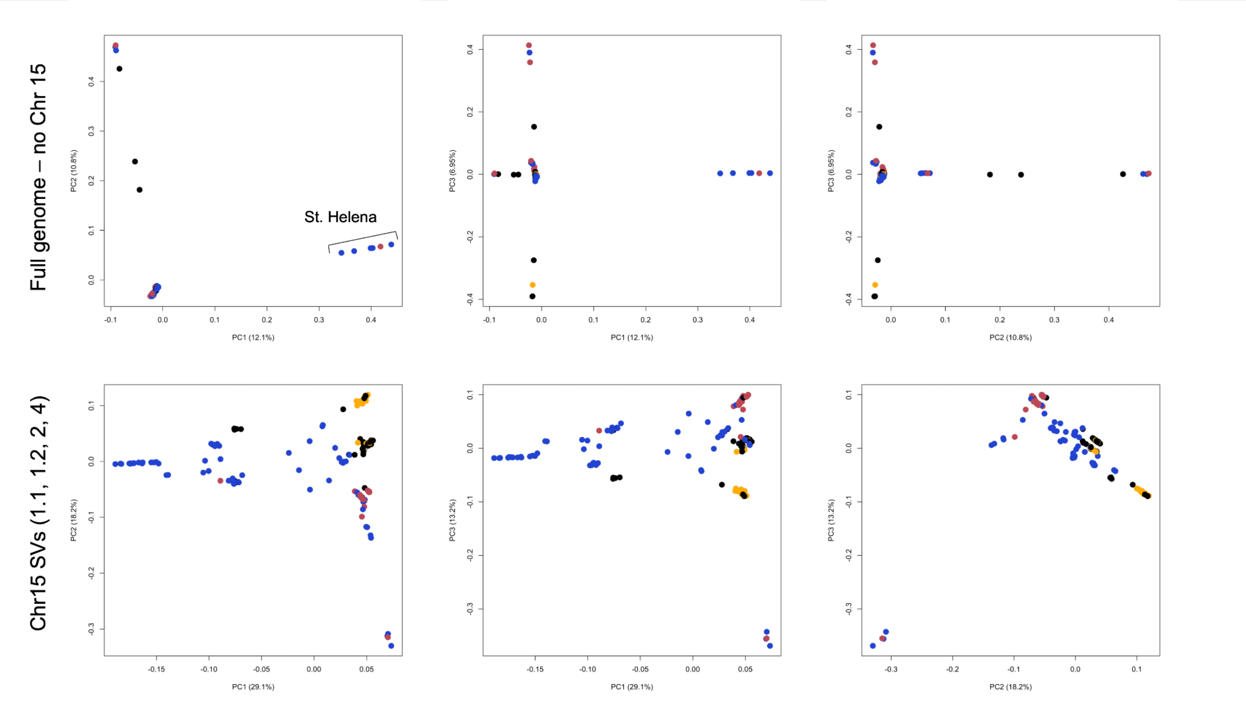


**Figure A. Principal Components Analysis.** PCAs representing genomic diversity across all 174 individuals using SNPs from all autosomes excluding chr15 (top) and from rearranged regions 1, 2, and 4 of chr15 [21] (bottom). Individuals are coloured by the morph to which they are assigned based on phenotype (see Fig 1A), chrysippus: red, klugii: yellow, orientis: blue, intermediate: black. The data underlying this Figure can be found in https://doi.org/10.5281/zenodo.14718778.

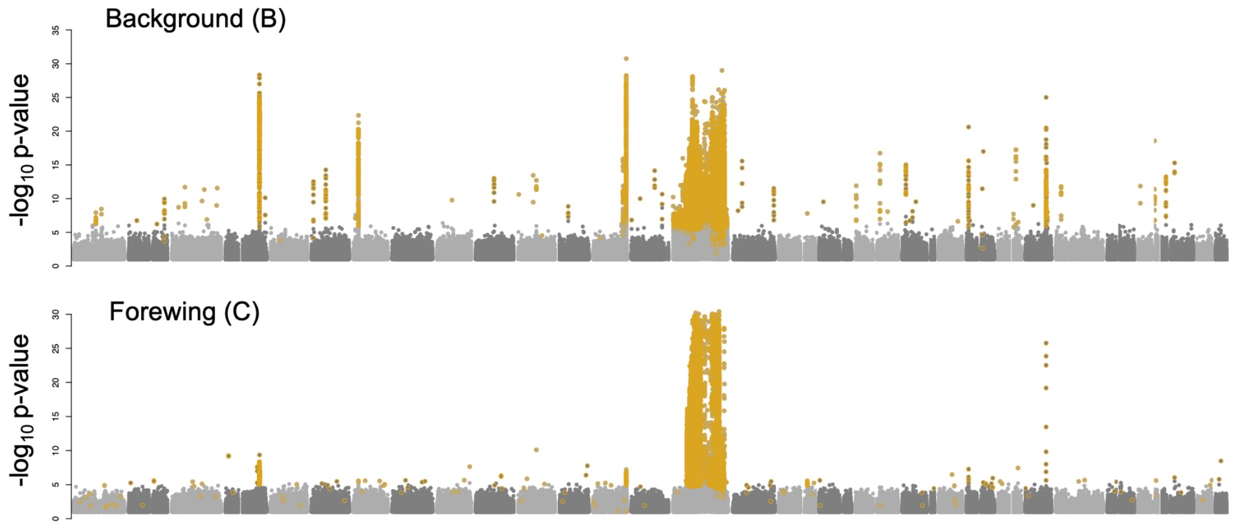


**Figure B. Association analysis using only hybrid zone individuals.** GWAS for the darkness of the wing background colour (top) and presence and absence of the forewing band (bottom) with only the 87 hybrid individuals we sampled. Significantly associated SNPs identified using a permutation test are coloured in yellow. The data underlying this Figure can be found in https://doi.org/10.5281/zenodo.14718778.

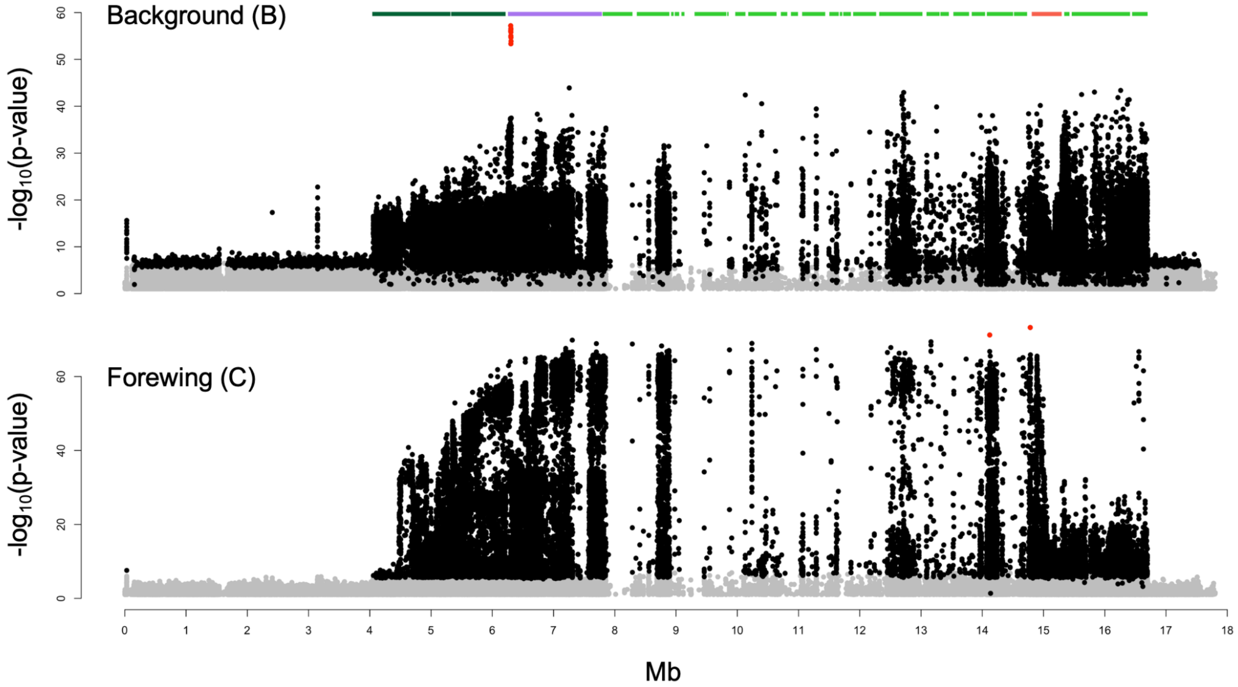


**Figure C. Association analysis results for chr15.** GWAS results for the darkness of the wing background colour (top) and presence and absence of the forewing band (bottom) across the full 172 individual dataset plotted across only chr15. The x-axis indicates chromosome position in megabases (Mb). The most strongly associated SNP peaks in each test ( -log10p-value > 50 for background and > 70 for forewing) are highlighted in red. The coloured bars above the plot indicate the locations of distinct rearranged tracts (or 'modules') described in [21]. The data underlying this Figure can be found in https://doi.org/10.5281/zenodo.14718778.

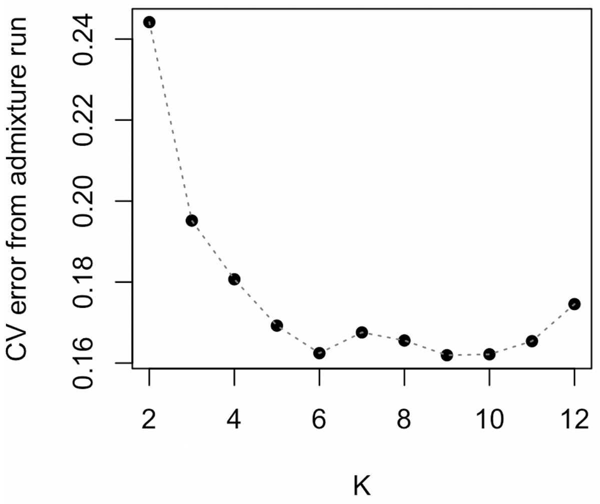


**Figure D Admixture cross validation (CV) error for the BC Supergene region (excluding the CNV region).** The data underlying this Figure can be found in https://doi.org/10.5281/zenodo.14718778.

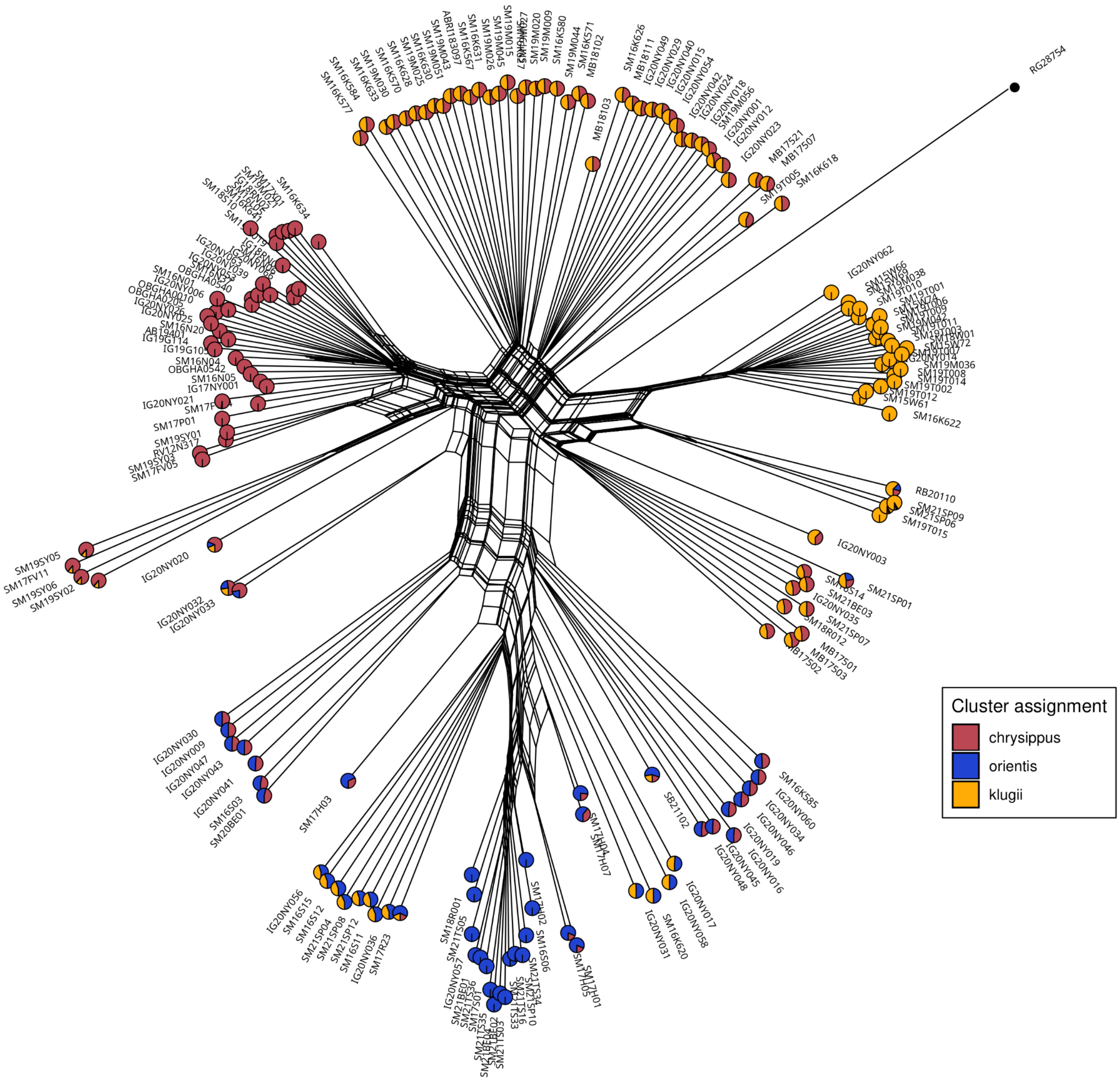


**Figure E. BC supergene clustering showing Admixture proportions with k=3.** Neighbor-Net network for unphased diploid genotypes across the BC supergene (excluding the CNV region). Each tip represents one diploid individual, and the network is constructed based on average pairwise genetic distances considering both haplotypes in each individual. We therefore expect 'heterozygous' individuals carrying two distinct haplotypes to be at intermediate positions in the network. Pie charts represent inferred ancestry components for each diploid individual from Admixture analysis with k=3 source populations. Population names assigned retrospectively based on known sample provenance. The data underlying this Figure can be found in https://doi.org/10.5281/zenodo.14718778.

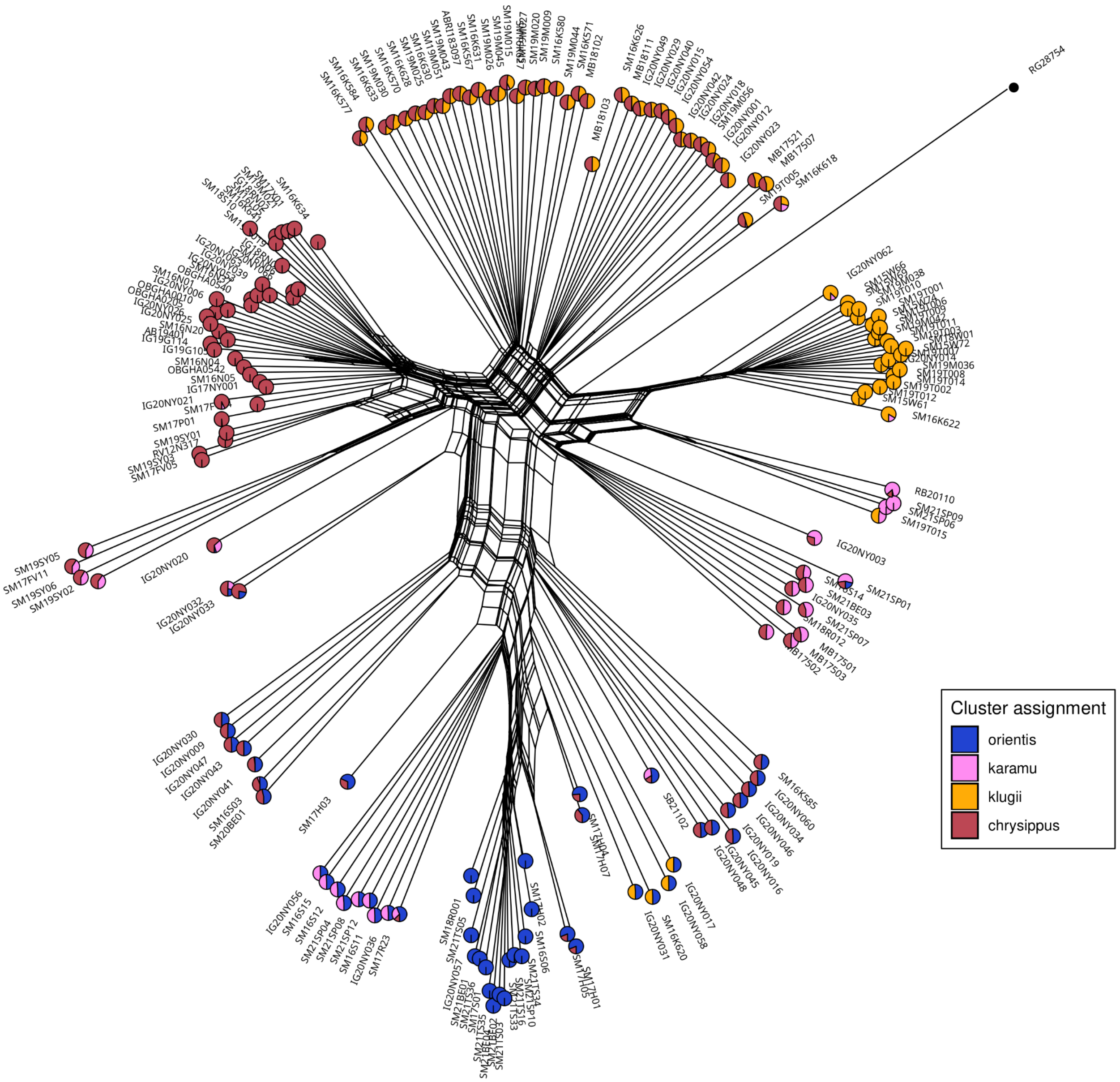


**Figure F. BC supergene clustering showing Admixture proportions with k=4.** Neighbor-Net network for unphased diploid genotypes across the BC supergene (excluding the CNV region). Each tip represents one diploid individual, and the network is constructed based on average pairwise genetic distances considering both haplotypes in each individual. We therefore expect 'heterozygous' individuals carrying two distinct haplotypes to be at intermediate positions in the network. Pie charts represent inferred ancestry components for each diploid individual from Admixture analysis with k=4 source populations. Population names assigned retrospectively based on known sample provenance. The data underlying this Figure can be found in https://doi.org/10.5281/zenodo.14718778.

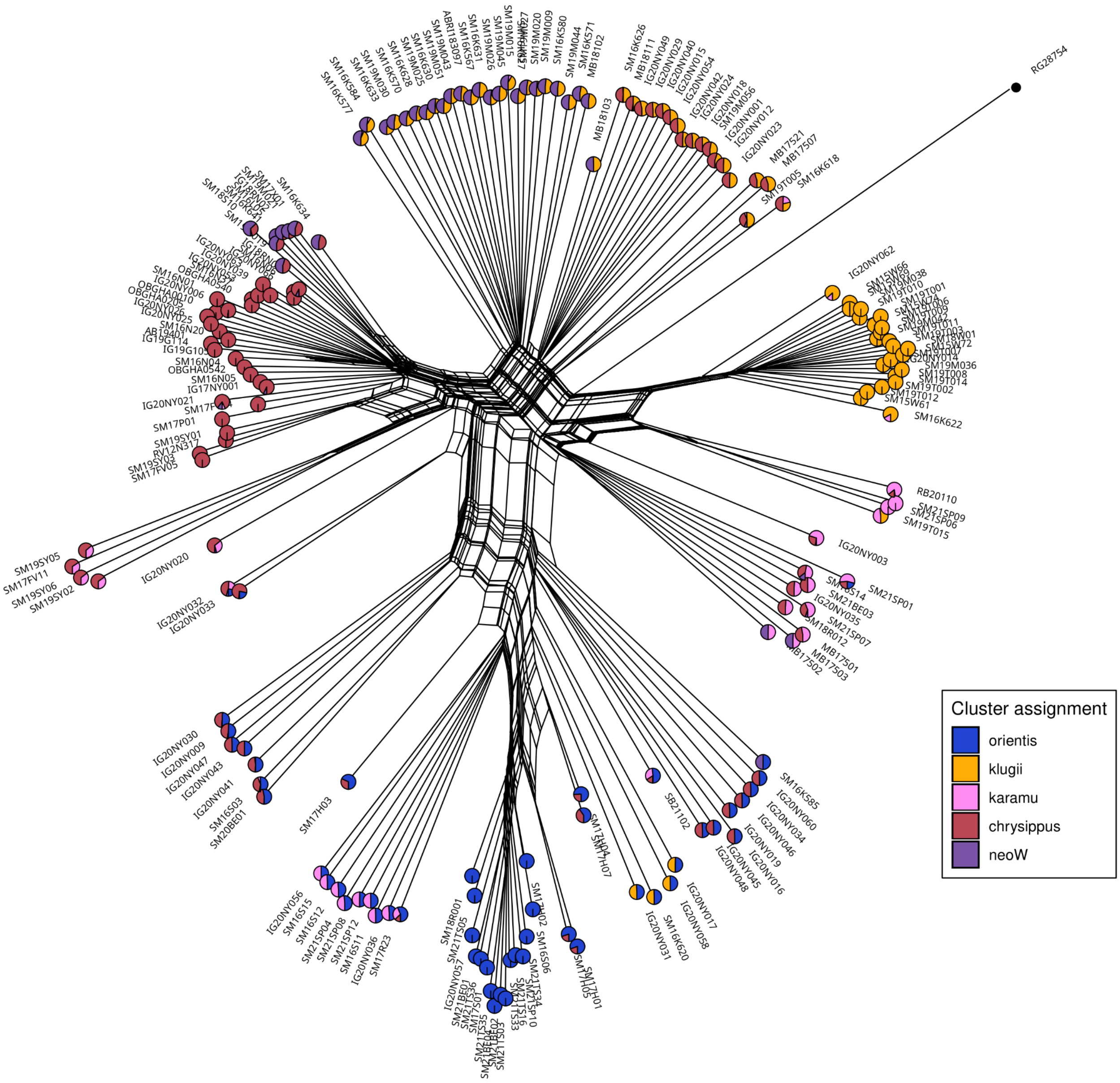


**Figure G. BC supergene clustering showing Admixture proportions with k=5.** Neighbor-Net network for unphased diploid genotypes across the BC supergene (excluding the CNV region). Each tip represents one diploid individual, and the network is constructed based on average pairwise genetic distances considering both haplotypes in each individual. We therefore expect 'heterozygous' individuals carrying two distinct haplotypes to be at intermediate positions in the network. Pie charts represent inferred ancestry components for each diploid individual from Admixture analysis with k=5 source populations. Population names assigned retrospectively based on known sample provenance. The data underlying this Figure can be found in https://doi.org/10.5281/zenodo.14718778.

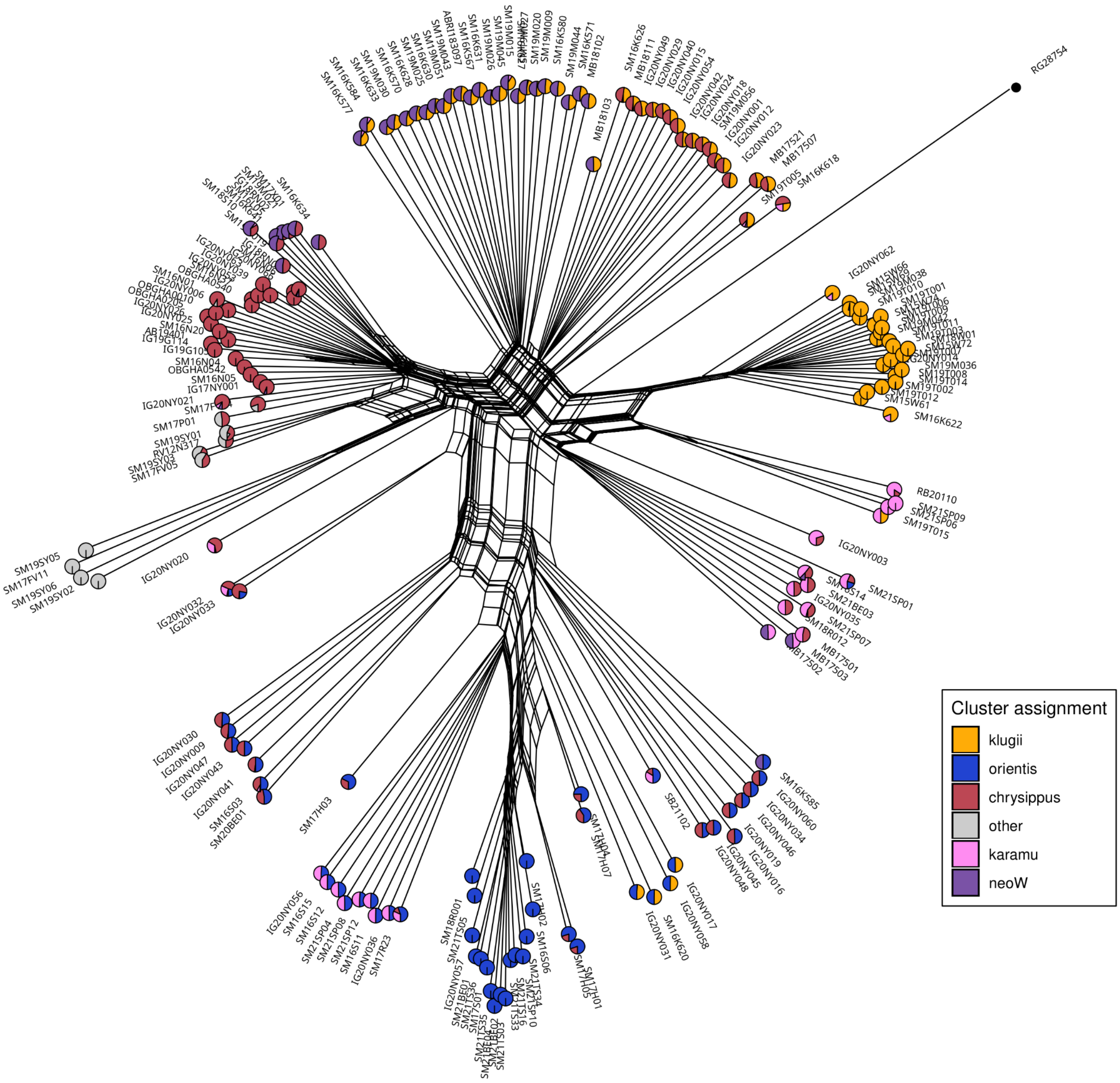


**Figure H. BC supergene clustering showing Admixture proportions with k=6.** Neighbor-Net network for unphased diploid genotypes across the BC supergene (excluding the CNV region). Each tip represents one diploid individual, and the network is constructed based on average pairwise genetic distances considering both haplotypes in each individual. We therefore expect 'heterozygous' individuals carrying two distinct haplotypes to be at intermediate positions in the network. Pie charts represent inferred ancestry components for each diploid individual from Admixture analysis with k=6 source populations. Population names assigned retrospectively based on known sample provenance. The data underlying this Figure can be found in https://doi.org/10.5281/zenodo.14718778.

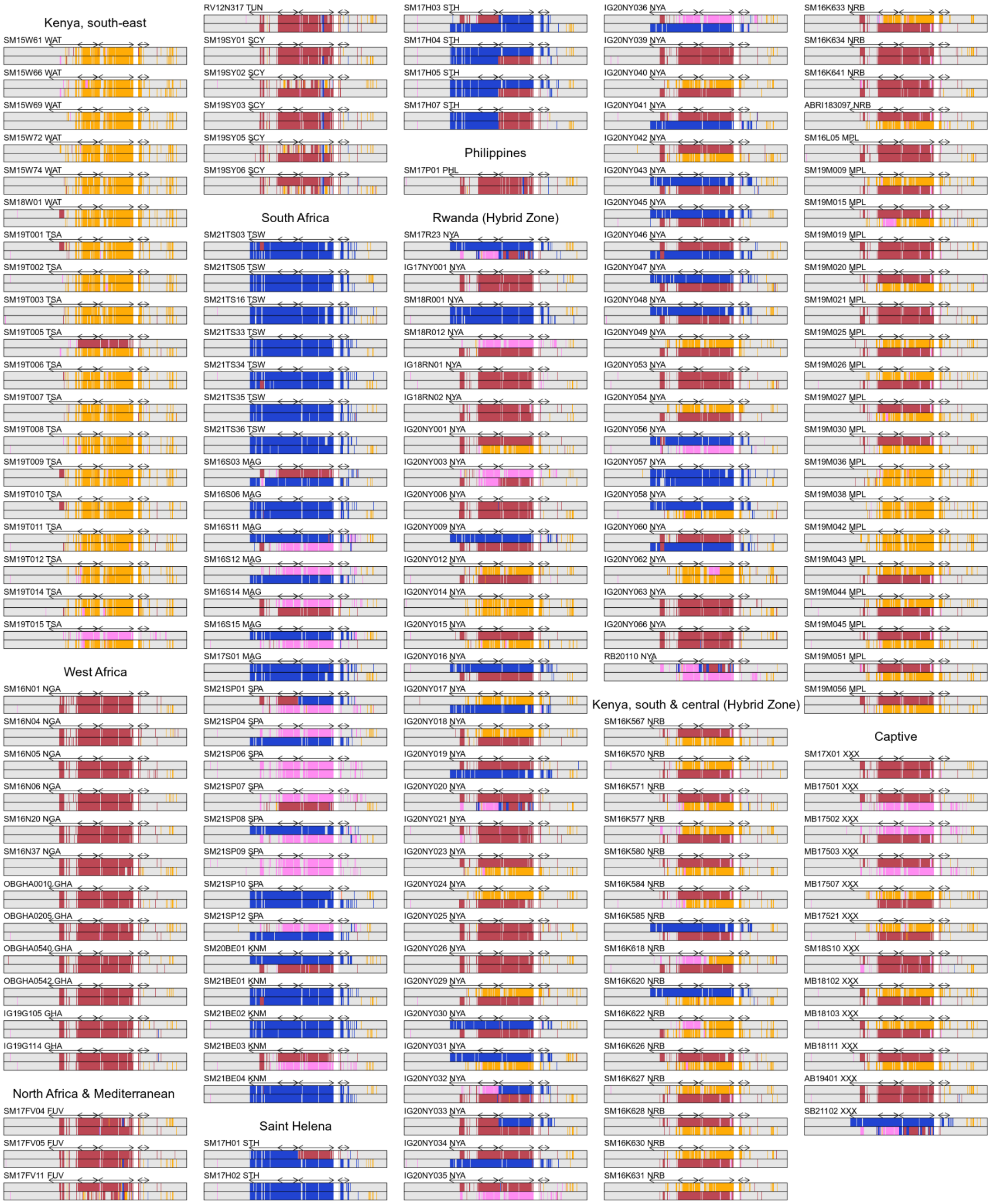


**Figure I. Ancestry painting of the BC supergene using the distPaint method.** For each individual, black outer boxes show the two haplotypes and vertical bars show ancestry assignments in windows of 200 SNPs with four source populations (yellow = klugii, red = chrysippus, blue = orients, pink = karamu, grey = unassigned [p > 0.01]). A subsection of chr15 from 2Mb to 17Mb is shown, with the CNV region replaced by a white gap. Arrows indicate the locations of inversions. The data underlying this Figure can be found in https://doi.org/10.5281/zenodo.14718778.

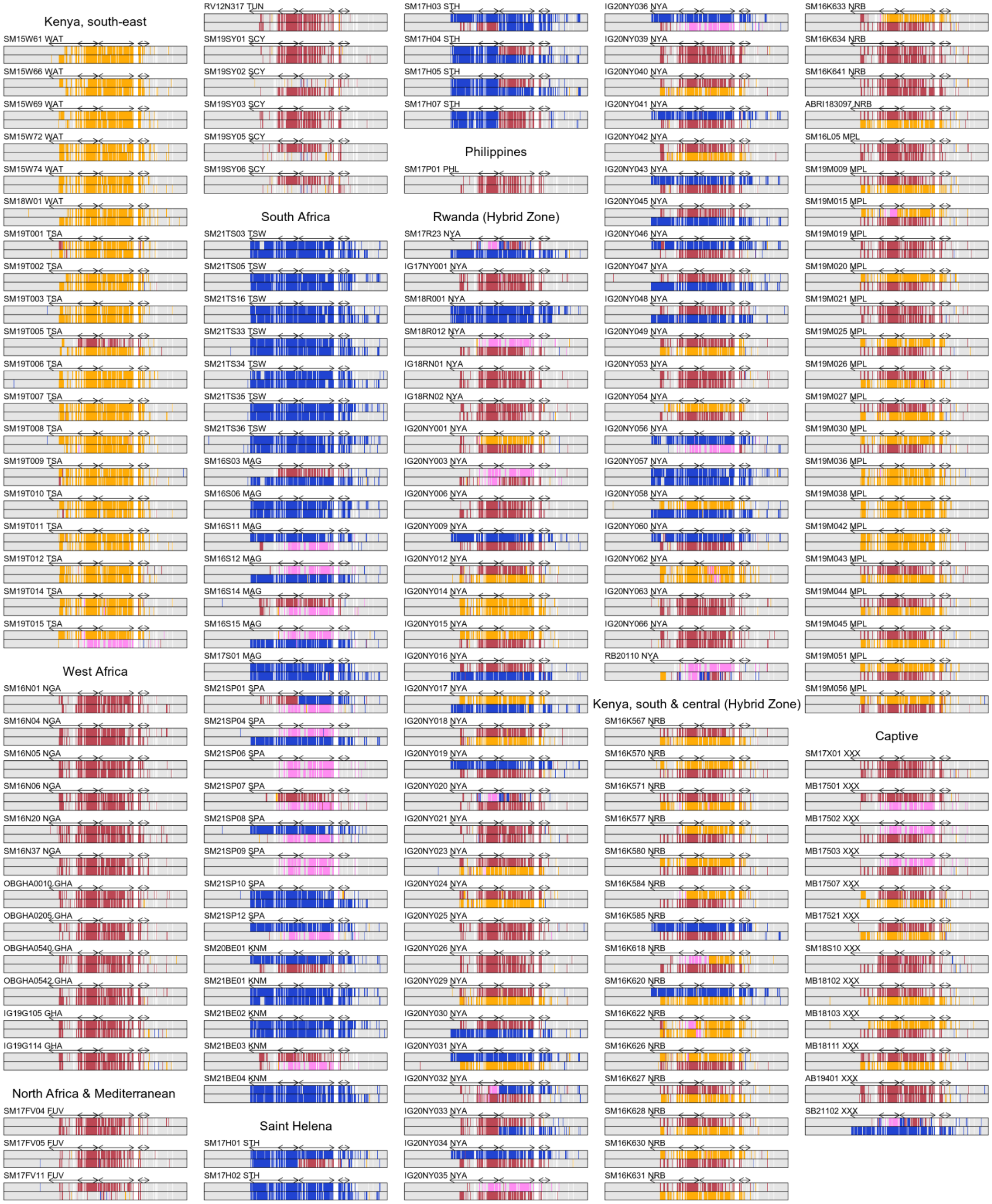


**Figure J. Ancestry painting of the BC supergene using Loter.** For each individual, black outer boxes show the two haplotypes and vertical bars show ancestry assignments for sets of SNPs according to the Loter hidden markov model with four source populations (yellow = klugii, red = chrysippus, blue = orients, pink = karamu, grey = unassigned). A subsection of chr15 from 2Mb to 17Mb is shown, with the CNV region replaced by a white gap. Arrows indicate the locations of inversions. The data underlying this Figure can be found in https://doi.org/10.5281/zenodo.14718778.

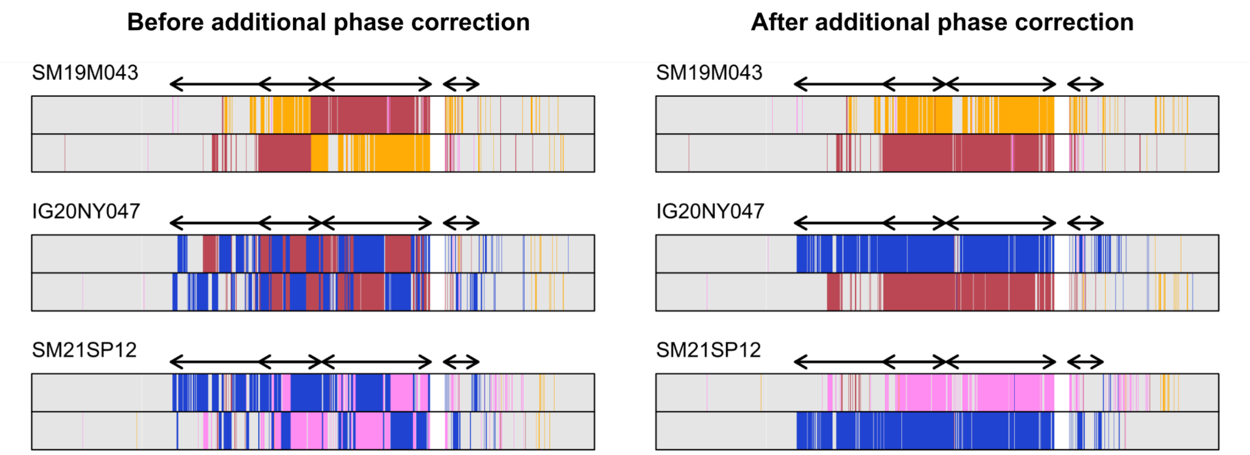


**Figure K. Examples of final phase correction step for ancestry painting.** All samples were phased using WhatsHap and Shapeit4 before ancestry painting analysis. Nevertheless, visual inspection of the results revealed multiple likely cases of phase switch errors, in which the two haplotypes of an individual appear as complementary mosaics of common haplotypes (left). Three representative individuals are shown. Black outer boxes show the two haplotypes in the individual, and vertical bars show ancestry assignments in windows of 200 SNPs (yellow = klugii, red = chrysippus, blue = orients, pink = karamu, grey = unassigned [p > 0/01]). Plots on the right show the same data after the additional phase correction algorithm has been run. Note that although only three individuals are shown, phase correction was performed using the complete set of 174 individuals. A subsection of chr15 from 2Mb to 17Mb is shown, with the CNV region replaced by a white box. Arrows indicate the locations of inversions. The data underlying this Figure can be found in https://doi.org/10.5281/zenodo.14718778.

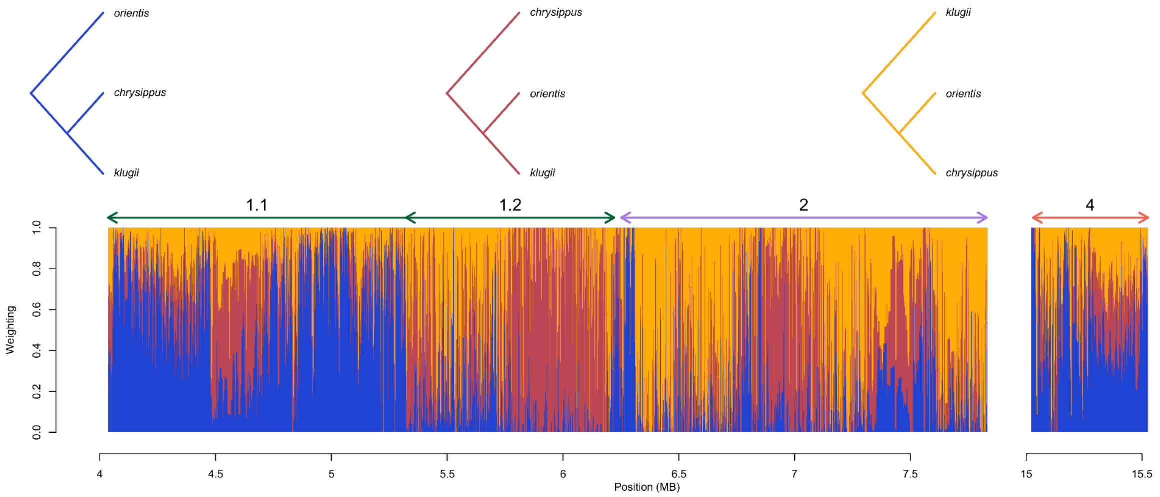


**Figure L. Topology weighting suggests historical gene flux in the BC supergene.** Coloured vertical bars show topology weights for three possible relationships between the prevalent haplotype groups of the BC supergene (trees shown above). Arrows and numbers along the top of the plot indicate the locations of inversions. Based on previous analyses [21], we expected that regions 1.1 and 4 (which are inverted only in orientis) would support the blue topology, which groups orientis apart from the other two. Likewise, region 1.2 (which is inverted in chrysippus) was expected to support the red topology, and region 2 (which is translocated in both chrysippus and orientis) was expected to support the yellow topology. Overall, these expected patterns are seen in the data, but the relationships between the haplotype groups also change at multiple points within each rearranged region, which is suggestive of historical gene flux through double-crossovers and subsequent breakdown of exchanged blocks through recombination. The data underlying this Figure can be found in https://doi.org/10.5281/zenodo.14718778.

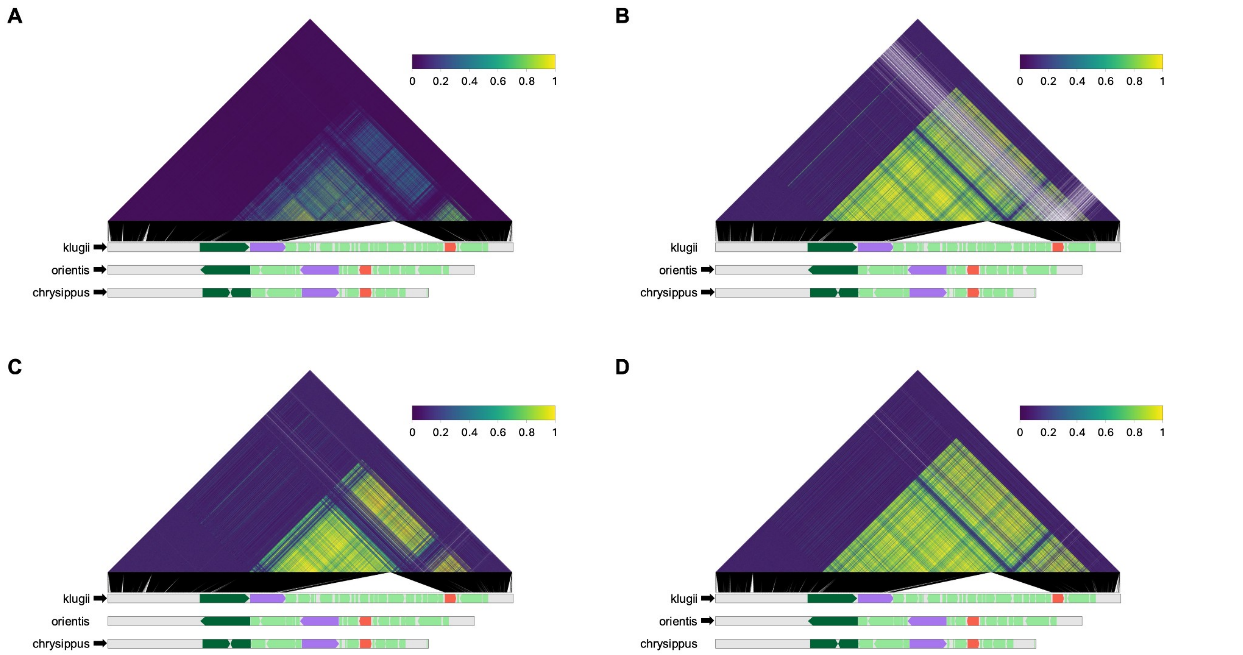


**Figure M. Linkage disequilibrium (LD) across chr15. A.** LD quantified using the $r^2$ squared metric between pairs of SNPs (minor allele frequency > 0.2) calculated across all 174 individuals. **B-D.** As in A, but using subsets of individuals representing two major haplotype groups in each case: B: chrysippus (NGA population) and orientis (TSW), C**:** klugii (WAT) and chrysippus (TSW), D: klugii (WAT) and orientis (TSW). The data underlying this Figure can be found in https://doi.org/10.5281/zenodo.14718778.

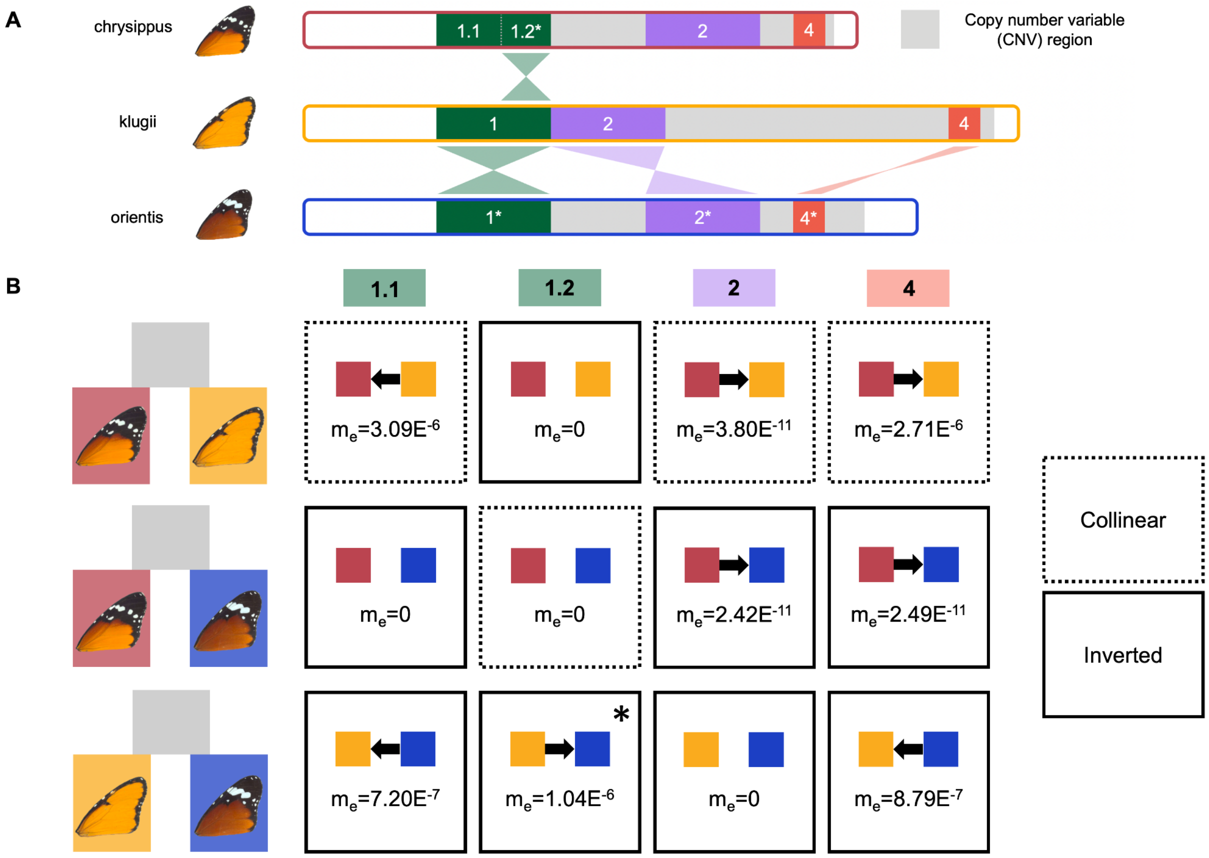


**Figure N. glMble analysis highlights gene flow between specific supergene module alleles. A.** A graphical representation of structural variation across the chrysippus, orientis, and klugii haplotypes replotted with data from [21] (regions with an inverted orientation relative to the ancestral state are indicated with *). **B.** Coalescent modelling between chrysippus and klugii individuals (top), chrysippus and orientis individuals (middle), and klugii and orientis individuals (bottom) highlighting which of the three model types, divergence only, divergence with gene flow from population A to B, and divergence with gene flow from B to A had the lowest composite likelihood for each region and comparison. Box borders indicate whether regions being compared are collinear (dotted line) or inverted (solid line) between morphs. * indicates the one case where the model failed to optimise. The values shown are provided in full in Table 2.
